# Integrating extracellular vesicle and circulating cell-free DNA analysis on a single plasma aliquot from breast cancer patients improves the detection of HER2 positivity

**DOI:** 10.1101/2023.03.02.530645

**Authors:** Vera Mugoni, Yari Ciani, Orsetta Quaini, Simone Tomasini, Michela Notarangelo, Federico Vannuccini, Alessia Marinelli, Elena Leonardi, Stefano Pontalti, Angela Martinelli, Daniele Rossetto, Isabella Pesce, Sheref S. Mansy, Mattia Barbareschi, Antonella Ferro, Orazio Caffo, Gerhardt Attard, Dolores Di Vizio, Vito Giuseppe D’Agostino, Caterina Nardella, Francesca Demichelis

## Abstract

**Background:** Multi-analyte liquid biopsies represents an emerging opportunity for non-invasive cancer assessment. We developed ONCE (ONe Aliquot for Circulating Elements), a novel multi-analytes liquid biopsy approach for the isolation of extracellular vesicles (EVs) and cell-free DNA (cfDNA) from a single aliquot of blood.

**Methods:** We assessed ONCE performance to classify HER2-positive early-stage breast cancer (BrCa) patients by combining RNA and DNA signals on n=64 healthy donors (HD) and non–metastatic BrCa patients. Specifically, we investigated EVs-derived RNA (EV-RNA) and cfDNA by next-generation sequencing (NGS) and by digital droplet PCR (ddPCR). Additionally, we utilized imaging flow cytometry to evaluate EVs as potential carriers of the HER2 protein.

**Results:** Western blot analysis and immunocapture assay revealed that EVs-enriched proteins were detected at similar levels among the HER2+ and HER2- subtypes. Sequencing of cfDNA and EV-RNA from HER2- and HER2+ patients demonstrated concordance with *in situ* molecular analyses of matched tissues. Combined analysis of the two circulating analytes by ddPCR showed increased sensitivity in *ERBB2/*HER2 detection compared to single nucleic acid components. Multi-analyte liquid biopsy prediction performance was comparable to tissue-based sequencing results from TCGA. Also, we observed HER2 protein on the surface of EVs isolated from the HER2+ BrCa plasma, thus corroborating the potential relevance of studying EVs as companion analyte to cfDNA.

**Conclusions:** This data confirms the relevance of combining cfDNA and EV-RNA analytes for cancer assessment and supports the ONCE approach as a valuable tool for multi-analytes liquid biopsies’ clinical implementation.

## Introduction

Liquid biopsies have emerged as a promising non-invasive tool to facilitate real-time detection and molecular profiling of primary and metastatic tumors [1]. Tests based on cell-free DNA (cfDNA) have been shown to hold diagnostic and prognostic significance in a range of cancer types [2], including BrCa [3, 4]. However, the sole analysis of cfDNA misses the information associated with gene expression, thereby limiting the potential utility of liquid biopsies.

Circulating materials actively investigated as transcriptomic resources include the RNA carried by circulating tumor cells (CTCs) and RNA circulating in the blood as free molecules or associated with extracellular vesicles (EVs). Specifically, the RNA cargo of extracellular vesicles (EV-RNA) contains a variety of high-quality RNA species, including protein-coding transcripts [5, 6] and offers the compelling advantage of being protected from degradation by the EV lipid bilayer and is overall more abundant than CTCs-associated RNA. Further, CTCs are extremely rare in early non-metastatic disease states [7]. Analysis by Next Generation Sequencing (NGS) or high-throughput technologies, such as quantitative real-time PCR (qRT-PCR), have already demonstrated the utility of EV-RNA for the detection of cancer–associated alterations in advanced BrCa [8, 9]. Furthermore, EVs represent an attractive circulating component for liquid biopsy tests as they act as carriers for multiple proteins, including cancer biomarkers such as the HER2 receptor in the context of BrCa [10–13].

We therefore sought to address the challenge of combining the analysis of circulating genomic and transcriptomic information by establishing a novel assay, implemented on a single plasma aliquot, that extracted useful information from both plasma DNA and EVs. Our assay is named ONCE (ONe Aliquot for Circulating Elements). It represents the first method to isolate EVs and cfDNA from a single plasma aliquot by combining existing validated procedures for EVs isolation and cfDNA extraction. We provide a proof-of-principle validation of EV-RNA and cfDNA isolated through ONCE as informative analytes for detecting the *ERBB2/HER*2 biomarker in a cohort of early-stage breast cancer (BrCa) patients with tissue-defined HER2 status.

Overall, our work implements a multimodal approach for multi-analytes liquid biopsies on a single aliquot of plasma, ultimately leading to improved detection sensitivity of cancer cargo at the early stages of the disease.

## Materials and Methods

### Experimental design and human sample collection

Blood samples from HDs were collected on a protocol approved by the University of Trento Ethics Committee (ID # 2017-010). BrCa patients’ plasma samples were prospectively collected on a protocol approved by the Ethics Committee of Santa Chiara Hospital in Trento (Rep.Int.12315 of July 24, 2017) with written informed consent. The experimental design was realized as described in **Supplementary Materials**.

### ONCE Protocol: combined isolation of EVs and cfDNA from human plasma

#### EV isolation by Charge-Based (CB) method and cfDNA extraction

The human plasma samples were filtered (Minisart NML syringe filters, pore size: 0.8 µm; Sartorius) and diluted with 1x Phosphate-Buffered Saline without calcium and magnesium buffer (Gibco) at 1:3 (v/v) ratio and processed for EV isolation by a charge – based isolation method as described before [14, 15]. The diluted plasma was recovered and processed for cfDNA isolation by QIAmp Circulating Nucleic Acid kit (Qiagen) [16].

#### EV isolation by ultracentrifugation (UC) and cell-free DNA (cfDNA) extraction

EVs were separated by UC on an Optima MAX-XP ultracentrifuge (Beckman Coulter) equipped with a TLA55 rotor. The plasma samples were filtered (Minisart NML syringe filters, pore size: 0.8 µm; Sartorius) and cleaned from cell debris by two serial centrifugation steps: 2000g for 10min and 10.000g for 20min. EVs were then pelleted at 100.000g for 70min, washed with 1mL of 1xPBS (Gibco), and re-pelleted at 100.000g for 70min as previously reported [17]. The liquid fractions (plasma and washing PBS) recovered from the UC steps were pooled in a separate clean tube and processed for cfDNA isolation as previously described [16].

### EV isolation by Size Exclusion Chromatography (SEC) (Izon qEV2 columns) and cfDNA Extraction

Plasma samples were filtered (Minisart NML syringe filters, pore size: 0.8 µm; Sartorius) and cleaned from debris by two serial centrifugation steps: 1.500g for 10 min followed by 10.000g for 10 min before loading on a pre-rinsed qEV2/70 nm column (Izon Science LTD, Cat. No. SP4). EVs were collected from 1^st^ to 5^th^ fractions by using an Izon Automatic Fraction Collector (AFC) and concentrated in Amicon Ultra 15 filtering units (Merck Millipore, Cat. No. UFC910024) by centrifugation at 3.000g for 20 min on a 5810R benchtop centrifuge (Eppendorf). The volumes eluted between the 6^th^ and the 20^th^ fractions were collected and processed for cfDNA isolation as previously described [16].

### Quantitation of EVs

The size and concentration of the EVs were quantitated using Tunable Resistive Pulse Sensing (TRPS, qNANO instrument, Izon Science) or Nanoparticle Tracking Analysis (NTA, NanoSight NS300, Malvern Panalytical). The method utilized for each data set is detailed in the figure legends and details reported in **Supplementary Materials**.

### Protein EV cargo profiling

EV proteins were extracted and analyzed by western blotting as previously reported [18] with few modifications as detailed in Supplementary Materials. EV surface marker analysis was performed by MACSPlex Exosome kit (Miltenyi Biotech; no. 130-108-813) following the manufacturer’s protocol tube for overnight capture on tubes. HER2 protein detection on EVs was performed by staining samples with HER2/ErbB2 (29D8) – PE-conjugated antibody or Isotype Control (Cell Signaling Technology) and CellMask plasma membrane stains (C10046, Life Technologies). HER2-positive EVs were identified on an Amnis ImageStream X MkII (Luminex).

### ddPCR assay

ddPCR assays performed on cfDNA and EV-RNA were from Bio-Rad Laboratories Inc (*ERBB2*: dHsaCP1000116; dHsaCPE5037554, EIF2C1: dHsaCP2500349, EEF2: dHsaCPE5050049) and PCR reactions were run on T1000 thermal cycler (Bio-Rad, Hercules, CA, USA). Analysis of ddPCR data was analyzed using QuantaSoft Software, version 1.7 (Bio-Rad Laboratories, Inc.). EV-RNA isolation was performed as detailed in **Supplementary Materials**.

### RNASeq on EV-RNA

EV-RNAs were processed with SMART-Seq® Stranded Kit (Takara Bio USA, Inc.) and libraries were sequenced on the Illumina HiSeq2500 platform by the Next Generation Sequencing Facility at the University of Trento (Italy). Reads were aligned against the human genome (hg38) using STAR [19]. Breast cancer tissue data [20] were downloaded from CBioPortal (https://www.cbioportal.org/), including immunohistochemistry evaluation of HER2.

### DNA whole-exome sequencing

Libraries were prepared with SeqCap EZ HyperCap Workflow version 2.3 (Roche). We utilized 20-50ng of cfDNA and 100ng of matched germline DNA (gDNA) that was sonicated to reduce the fragment size to 180-220bp (Covaris M220). Libraries were sequenced on the Illumina HiSeq2500 platform by the Next Generation Sequencing Facility at the University of Trento (Italy). CNVkit [21] was used for copy number aberration (CNA) detection via segmentation, and Log2 values of cfDNA over control were corrected for purity and ploidy using ClonetV2 [22]. Extended procedure and details are presented in **Supplementary Materials**.

### HER2 positivity classification of BrCa study patients

We used ddPCR data for all liquid biopsy samples (the highest DNA and RNA ddPCR values of healthy donor samples were used as lower thresholds for DNA and RNA, respectively) and sequencing data for the TCGA tissue samples (DNA amplification threshold was set at 2.6 copies; RNA levels for samples without DNA amplification were considered and the 75% percentile of the distribution was set as lower threshold). For each sample, we then considered the following classes of HER2+ positivity; *Only RNA*: RNA but not DNA signal is higher than the corresponding threshold; *Only DNA*: DNA but not RNA data is higher than the corresponding threshold; *Combo AND*: both DNA and RNA data are above the corresponding thresholds; *Combo OR*: any of DNA or RNA data is higher than the corresponding threshold. The classification performance was estimated based on precision, recall and F1 score as described in **Supplementary Materials**.

### Statistical Analysis

Statistical analysis was performed using GraphPad Prism version 8.0 software (www.graphpad.com) and R software (www.R-project.org). One-way ANOVA or t-test were used as appropriate. Data visualizations were performed using the packages ggplot2 (https://ggplot2.tidyverse.org) and complexHeatmap.

Additional methods and procedures are described on **Supplementary Materials.**

## Results

### ONCE combines EV isolation with cfDNA extraction from a single plasma aliquot

Previously reported procedures for isolating cfDNA and EVs require different aliquots of body fluids to purify each circulating component [23, 24]. We hypothesized that the concomitant isolation of two diverse analytes (cfDNA and EVs) from the same aliquot of a body fluid might be feasible, reduce the amount of valuable blood required for testing, and inherently avoid inter-sample variability that might affect the sensitivity in detecting relevant cancer information. To this purpose, we developed and tested the ONCE approach as a novel combination of serial procedures for isolating EVs and cfDNA from a single plasma aliquot (workflow in Fig. 1a). To assess its applicability with a range of EV isolation methods, we tested multiple approaches. Specifically, we collected whole blood from a cohort of HDs (n=20) into EDTA tubes (commonly used for EVs isolation) and separated the plasma fraction. We then divided the collected plasma into three identical aliquots and processed each one of them with a diverse EV isolation method: 1) a charge–based (CB) method [15, 25], previously used for analyses on EVs isolated from cancer patient plasma [26]; 2) ultracentrifugation (UC), the most commonly utilized EV isolation method [17]; 3) size-exclusion chromatography (SEC) on qEV/70nm columns associated with an automatic fraction collector system certified as a medical device (IZON Science LTD) [27].

**Figure 1:**
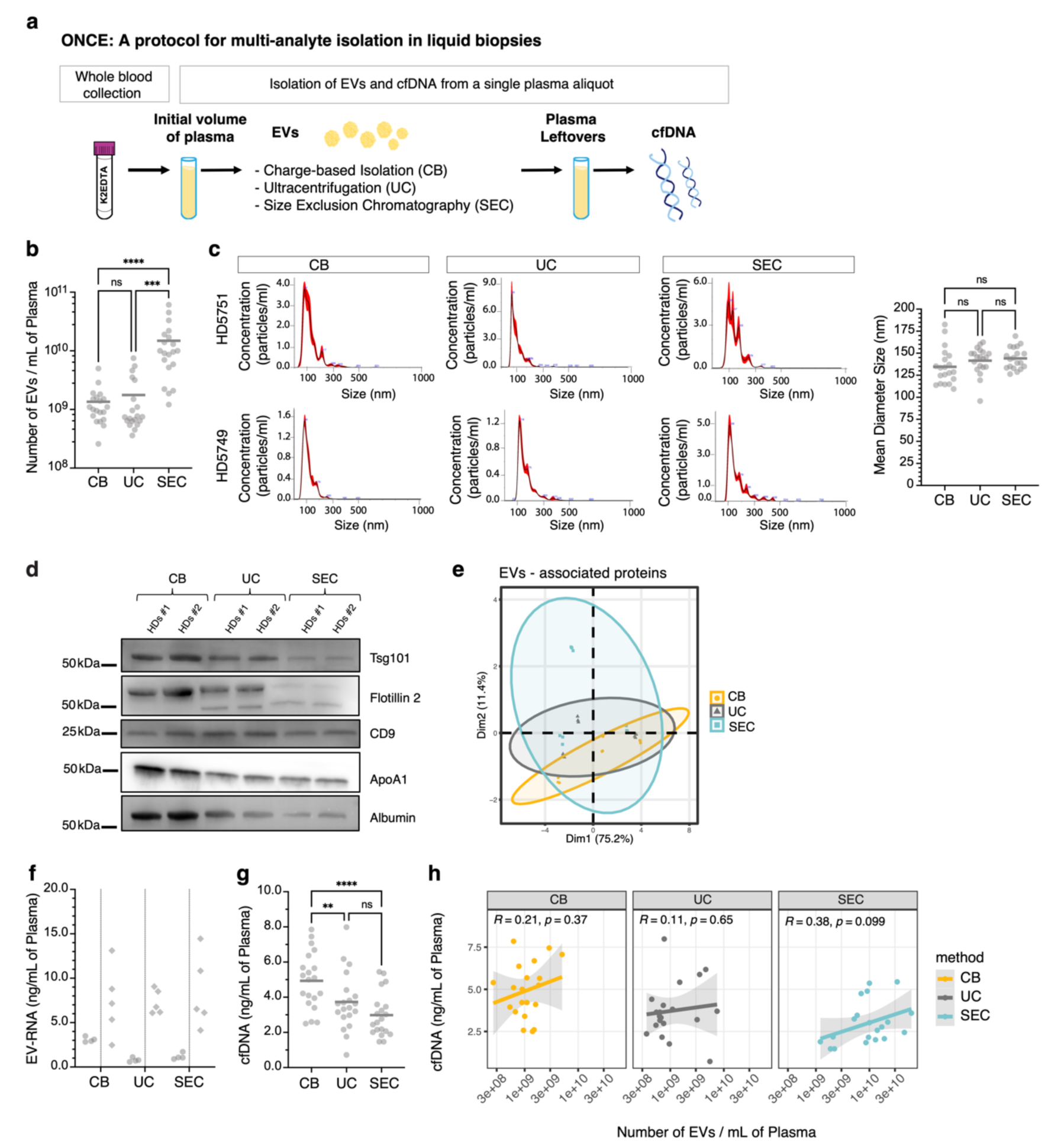
ONCE combines diverse EV isolation methods and cfDNA extraction from a single plasma aliquot. **a.** Scheme of ONCE (**<u>ON</u>**e Aliquot for **<u>C</u>**irculating **<u>E</u>**lements) protocol. Starting from a single plasma aliquot, we propose the ONCE protocol as a methodological approach for sequentially isolating two analytes circulating in the blood: the EVs and the cfDNA. We evaluated three diverse methods for EV isolation: CB, UC, and SEC. After EVs isolation, plasma leftovers are utilized for cfDNA extraction. **b.** Quantification of EVs by NTA measurements. EVs were isolated from a cohort of HDs (n=20). Three plasma aliquots were collected from each HDs and EVs were isolated from each aliquot using one of the three specific methods: CB, UC, or SEC. Compared to CB and UC, the SEC method resulted in higher nanoparticle recovery. All plasma for EVs isolation was derived from blood collected into K2EDTA tubes. ns: not significant differences; *** p= 0.0001, **** p< 0.0001 by 2way ANOVA. **c.** Representative profiles of EV samples quantitated in **panel b**. EVs were isolated through ONCE using one of the three specific methods: CB, UC, and SEC. EVs were analyzed by NTA (Nanoparticle Tracking Analysis), and quantitation of EVs mean diameter size indicates that all isolated samples correspond to small EVs. NTA profiles are from n=2 EVs samples of representative healthy individuals (HD5751 and HD5749) and are shown as a merge of n=5 acquired measurements (videos). The mean diameter size was calculated by NanoSight NTA software v3.2. n.s.: not significant differences by One – way Anova. **d.** Western Blot assay showing the EV-enriched proteins Tsg101, Flotillin 2, CD9, and the contaminant proteins Apolipoprotein A1 (ApoA1) and Albumin in representative n=2 pool of EV samples from HDs. Plasma from the same individuals was divided into equal aliquots, and EVs were isolated from plasma by CB, UC, or SEC. **e.** Principal component analysis (PCA) on signals derived from 37 exosomal surface epitopes detected on EV samples of n=3 HDs (technical replicates n=3). EVs were isolated by CB, UC, or SEC, and EV surface markers were profiled by multiplex bead-based flow cytometry assay (MACSPlex exosomal kit; Miltenyi Biotec). Each ellipse groups the exosomal epitopes’ expression profile of EVs isolated using one specific method. EVs isolated from the same individuals by diverse methods show partial overlap. **f.** Quantification of EV-RNA extracted from EVs after isolation. EV-RNA was quantitated by Agilent RNA pico assay (left) or small RNA assay Kit (right). EV-RNA samples were derived from n=9 HDs. The plasma of each HDs was divided into three aliquots and each plasma aliquot was processed for EVs isolation by using a dedicated method (CB, UC, SEC). Separated EVs were later processed for RNA extraction as detailed in methods. **g.** Quantification of cfDNA by Qubit dsDNA HS Assay (Thermo Fisher Scientific), extracted from plasma leftovers collected after EVs isolation. Data are from n=20 HDs. cfDNA was extracted from plasma leftovers, collected after EVs were isolated with a dedicated method (CB, UC, SEC) as shown in **panel a**. h. Scatter plots correlating the amount of recovered cfDNA and EVs from each HDs (n=20) across the three EV isolation methods (CB, UC, SEC). No significant positive or negative correlation is derived. R and p-values are obtained from linear model (lm function in R). **cfDN**A: cell free DNA; **EV**: Extracellular Vesicles; **EV-RNA**: RNA extracted from EVs. **HDs**: Healthy Donors; **CB**: Charge – Based Isolation Method**, UC**: Ultracentrifugation; **SEC**: Size – Exclusion Chromatography. See also associated Supplementary Fig. 1 and Supplementary Fig. 2

After EV isolation, we recovered the plasma leftovers and processed them for cfDNA extraction through a silica-based membrane technology [16, 23] (**Fig. 1a**).

First, we compared the efficiency of the three considered methods for EV isolation by performing nanoparticle tracking analysis (NTA) measurements. While the SEC method resulted in the highest nanoparticle recovery efficiency (**Fig. 1b**), the profiles and the mean diameter of the EV samples were similar across the methods, thereby suggesting that all three methods allow efficient isolation of small EVs (**Fig. 1c**).

Immunoblot detection of protein markers commonly associated with exosomes (Tsg101, Flotillin-2, CD9) as well as commonly co-isolated contaminants (Apolipoprotein A1 and Albumin) revealed a distinctive expression profile across the EV samples obtained from the same individuals subjected to the diverse isolation methods (**Fig. 1d**). To confirm the distinctiveness of EVs obtained by CB, UC or SEC, we performed a comparison on the detection levels of a panel of 37 exosome surface epitopes by MACSPlex immunocapture assay [28]. PCA analysis showed partial overlap among the samples isolated with the different methods suggesting that the expression levels of the analyzed epitopes are only partially concordant upon EVs isolation through diverse methods (**Fig. 1e**).

We then evaluated the recovery of nucleic acids associated with EVs isolated from HDs using the three diverse methods. Starting from small EVs isolated from an aliquot of plasma (1.8mL), we extracted a minimal amount of DNA (less than 1ng of DNA per mL of plasma) across the three EV isolation methods (**Supplementary Fig. 1a**), thereby confirming previous data reporting DNA mainly carried by large EVs [29]. By using the same volume of plasma (1.8mL) for EVs isolation by CB, UC, or SEC, we were able to extract a good amount of RNA (to 10 - 15 ng of RNA per mL of plasma) (**Fig. 1f**) with comparable fragment size length across the tested EV isolation methods (**Supplementary Fig. 1b**).

Once we assessed the differences in recovered EV-RNA among CB, UC, and SEC, we analyzed the relative plasma leftovers (EV-depleted plasmas) for the subsequent extraction of cfDNA. EV-depleted plasmas obtained after the CB-mediated EV isolation method resulted in the highest cfDNA recovery when compared to EV-depleted plasmas collected after UC and SEC processing (**Fig. 1g**); in contrast, the size length of the extracted cfDNA fragments was comparable among the three **(Supplementary Fig. 1c**). Thus, the CB method resulted as the most convenient approach for combining the extraction of cfDNA with the isolation of EVs. Also, we assessed that the efficiency of cfDNA recovery was not directly related to EV extraction’s efficiency. We did not observe a significant correlation between the amount of recovered cfDNA and the number of isolated EVs across the three methods for EV isolation (CB, UC, and SEC), thus suggesting that the differences in cfDNA recovery between the three methods are independent of the efficiency of EV isolation (**Fig. 1h**).

To further assess whether the efficiency of cfDNA extraction from EV-depleted plasma was affected by the prior isolation of EVs, we checked if the levels of cfDNA extracted by performing ONCE with the CB isolation method were comparable with the cfDNA levels obtained without any plasma processing steps for the EV isolation (I CONTROL and II CONTROL) (**Supplementary Fig. 2a**). To this purpose, we collected blood from n=20 HDs. Two types of tubes commonly utilized for liquid biopsies, i.e., K2EDTA and BCT Streck tubes, were used. Each volume of the separated plasma sample was split into three aliquots (1.5ml/each). The three aliquots of plasma were then processed, each with one of the three following protocols (**Supplementary Fig. 2a**):

i) ONCE Protocol, the aliquot is diluted in PBS1X and incubated with EV-capture beads for EV isolation; the diluted EV-depleted plasma leftovers are then utilized for cfDNA extraction.
ii) I CONTROL protocol, the aliquot is processed to purify cfDNA according to the protocol commonly utilized for cfDNA isolation [30];
iii) II CONTROL protocol, the aliquot is first diluted in PBS1X (as in *i*) and then processed to purify cfDNA, as in *ii*, to check whether the sole dilution of the plasma (performed to reduce plasma viscosity) interferes with the efficiency of cfDNA recovery.

By comparing the ONCE protocol with controls (protocols I-CTRL and II-CTRL), no significant differences were detected in the amount or in the size length of cfDNA recovered, regardless of the blood tube type and the relevant associated preservatives (**Supplementary Figure 2b-g**).

Altogether, our data indicate that the CB method for EV isolation within the framework of the ONCE approach allows for the combined isolation of cfDNA and EVs and ensures the preservation of the isolated DNA from a single plasma aliquot.

In light of these results, for the subsequent experiments, we opted for the CB-mediated EV isolation method in the framework of ONCE as this method combines an EVs recovery (in terms of the EVs number isolated per mL of plasma) comparable to UC with an optimal cfDNA yield but is more rapid and easily scalable.

### Processing of liquid biopsies from early-stage BrCa patients for serial isolation of EVs and cfDNA

To assess the potential of detecting cancer-derived cargo in liquid biopsies from cancer patients, we tested ONCE in a cohort of early-stage BrCa patients (n=44) prospectively enrolled between 2017 and 2019 at the Santa Chiara hospital in Trento (Italy) and subjected to neoadjuvant chemotherapy (NAC) (**Table 1**). Specifically, we sought to test the ability to assess HER2 status in patients’ circulation by querying multiple analytes. To this end, baseline plasmas from single aliquots of blood collected into a K2EDTA tube were utilized to extract cfDNA after EV isolation by the CB method (**Fig. 2a**).

**Figure 2:**
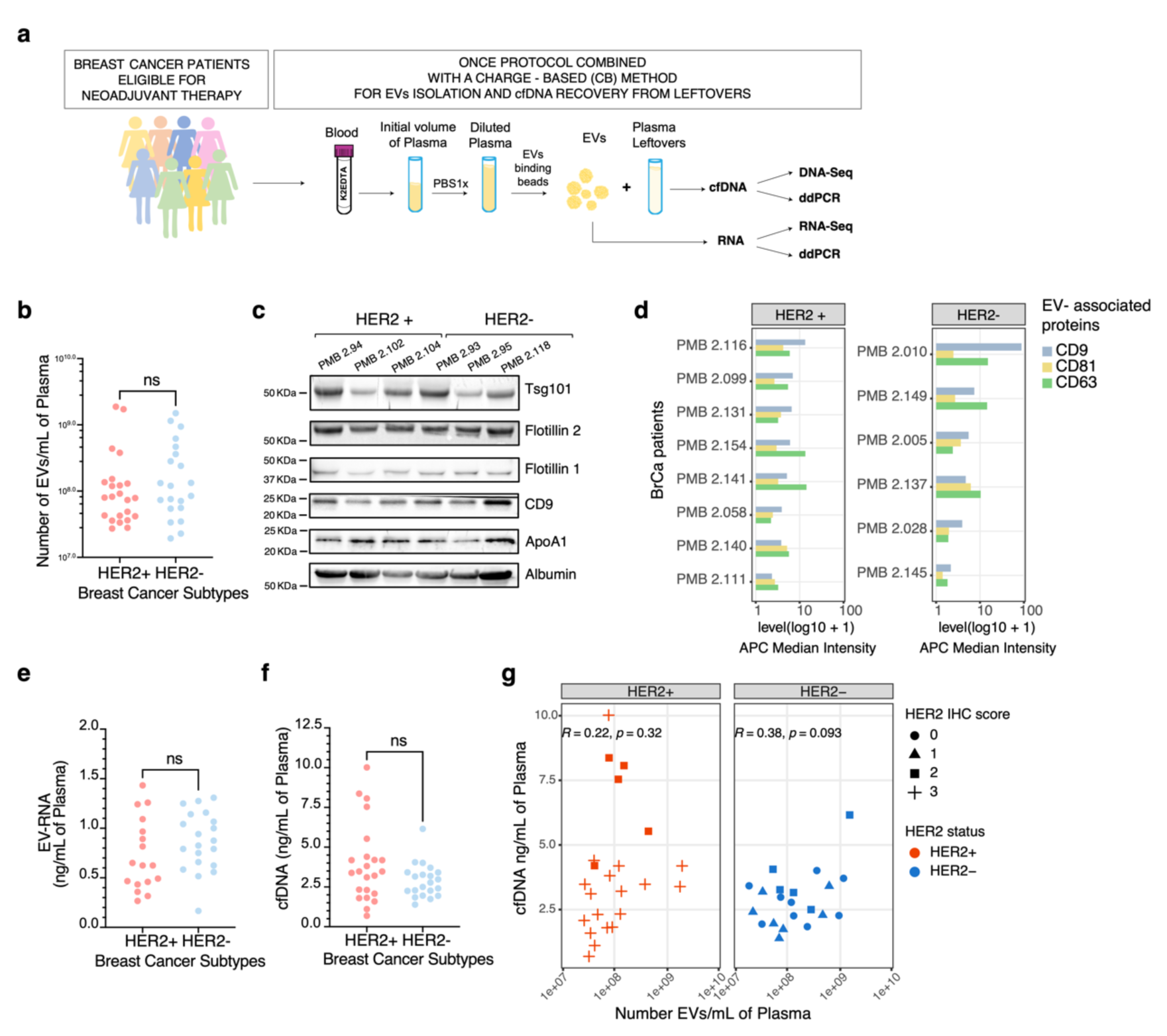
EVs and cfDNA are concomitantly and efficiently extracted from early-stage BrCa liquid biopsies. **a.** Scheme of ONCE (**<u>ON</u>**e Aliquot for **<u>C</u>**irculating **<u>E</u>**lements) protocol for multi-analyte liquid biopsies in BrCa. An aliquot of whole blood was collected from n=44 early-stage non metastatic BrCa patients into K2EDTA tubes and the extracted plasma (1.5ml) was processed by ONCE protocol combined with the CB method for the isolation of two diverse circulating analytes: EVs and cfDNA. EVs and cfDNA are sources of circulating nucleic acids for investigating tumor biomarkers by RNA-Seq/DNA-Seq and /or ultra-sensitive ddPCR. The dilution of plasma with PBS1x is an essential step to reduce plasma viscosity, thereby facilitating the binding of the beads to the EVs and guaranteeing the performance of the CB method. **b.** Quantification of EVs by TRPS (Tunable Resistive Pulse Sensing) measurements. EVs were isolated from early-stage breast cancer patients (n=44) of 2 different subtypes: HER2+ and HER2-. n.s.: not significant differences by t-test. **c.** Western Blot assay showing the EV-enriched proteins Tsg101, Flotillin 2, Flotillin 1, CD9 and the contaminant proteins Apolipoprotein A1 (ApoA1) and Albumin in representative n=3 HER2+ and n=3HER2- EV samples isolated from BrCa patients (PMB). EVs were isolated by CB isolation method as shown on **panel a**. **d.** Detection of CD8, CD63, and CD81 tetraspanins by MACSPlex exosome assay on EVs isolated by CB method throughout ONCE protocol. X axis plots the detection signal (Median Intensity level on APC channel) in log10+1 scale. Y axis reports BrCa patient code. Data are from n= 8 HER2+ and n=6 HER2- BrCa patients. **e.** Quantification of EV-RNA extracted from EVs after isolation by CB method. EV-RNA was quantitated by Agilent RNA pico assay. EV-RNA samples were derived from n=17 HER2+ and n=20 HER2- early-stage BrCa patients. n.s.: not significant differences by t-test. **f.** Quantification of cfDNA by Qubit dsDNA HS Assay (Thermo Fisher Scientific). cfDNA was extracted from plasma leftovers, after EVs enrichment, according with ONCE protocol. Data are from n=23 HER2+ and n=21 HER2- early-stage BrCa. n.s.: not significant differences by One–way Anova. **g.** Scatter plot of levels of cfDNA (ng/mL of plasma) and EVs (number of EVs/ mL of plasma) recovered by ONCE from n=44 BrCa patients. Data are stratified by HER2 status (HER2+ vs HER2-) based on histopathological classification. The IHC score, derived from tissue biopsies, is shown with symbols. **cfDN**A: cell free DNA; **EVs**: Extracellular Vesicles; **EV-RNA**: RNA extracted from EVs. **BrCa**: Breast Cancer; **HER2+**: HER2 positive, **HER2-**: HER2 negative; **IHC**: Immunohistochemistry. See also Supplementary Fig. 3.

**Table 1.** Demographic table of breast cancer patients’ study cohort

| Characteristic | Patients (N=56) |
| --- | --- |
| **Age at baseline** |  |
| Median age (range) - yr | 52.9 (28-77) |
| 40 years - no. of patients (%) | 4 (7.1%) |
| > 40 years - no. of patients (%) | 52 (92.8%) |
| **Subtype Classification - no. of patients (%)** |  |
| HER2 positive: HER2+ | 17 (30.3%) |
| Luminal B (HER2+): ER+ a/o PR+ HER2+ | 10 (17.8%) |
| Triple Negative: ER- PR- HER2- | 16 (28.5%) |
| Luminal B (HER2-): ER+ a/o PR+ HER2- | 13 (23.2%) |
| **Histological Subtype - no. of patients (%)** |  |
| Ductal | 51 (91.0%) |
| Lobular | 2 (3.6%) |
| Infiltrating NST | 3 (5.4%) |
| **HER2 Expression Status (IHC; score) - no. of patients (%)** |  |
| 0, 1+ (negative) | 17 (30.3%) |
| 2+ (ambiguous) | 12 (21.4%) |
| 3+ (positive) | 26 (46.4%) |
| **HER2 Amplification Status (FISH)\* - no. of patients (%)** |  |
| Positive | 9 (16.0%) |
| Negative | 7 (12.5%) |
| Not performed* | 40 (71.4%) |
| **Metastasis (M; Pathological stage) - no. of patients (%)** |  |
| M0 | 56 (100%) |
| **Clinical Cancer Stage - no. of patients (%)** |  |
| Stage IA-IB | 6 (10.7%) |
| Stage IIA | 19 (33.9%) |
| Stage IIB | 16 (28.6%) |
| Stage IIIA | 6 (10.7%) |
| Stage IIIB | 3 (5.3%) |
| Stage IIIC | 3 (5.3%) |
| n.a. | 3 (5.3%) |
| *According to ASCO-CAP guidelines (Wolff AC, Hammond MEH, Allison KH, Harvey BE, Mangu PB, Bartlett JMS, et al. Human Epidermal Growth Factor Receptor 2 Testing in Breast Cancer: American Society of Clinical Oncology/College of American Pathologists Clinical Practice Guideline Focused Update. J Clin Oncol 2018;36(20):2105-22 doi 10.1200/JCO.2018.77.8738). |  |

We first quantified the isolated EVs by tunable resistive pulse sensing (TRPS) technology. We observed no significant differences in EV concentration between HER2+ and HER2- subtypes (**Fig. 2b**) (mean range across subtypes: 2.40E+08 and 3.10E+08 EVs/mL of plasma, respectively). By western blot analysis and MACSPlex immunocapture assay, we confirmed the expression of EVs-enriched proteins, including Tsg101, Flotillin 1/2, and tetraspanins (CD9, CD63, CD81) at similar levels among a subset of the HER2+ and HER2- samples (**Fig. 2c,d**).

Next, to evaluate the EV-associated nucleic acid cargo, we extracted DNA and RNA from these early-stage BrCa patients’ EVs (EV-DNA and EV-RNA). A minimal amount of EV-DNA (mean: 0.0140 ng of DNA/ mL of plasma) was obtained from patient plasma (**Supplementary Fig. 3a**). In contrast, higher DNA recovery was obtained from synthetic liposomes containing artificial DNA and from EVs of metastatic BrCa patient plasmas, here included as internal controls (**Supplementary Figure 3a-c**). Regarding the EV-RNA, we consistently extracted comparable amounts among HER2+ (mean: 0.729 ng/mL of plasma) and HER2- samples (mean:0.866 ng/mL of plasma) (**Fig. 2e** and **Supplementary Fig. 3d**).

Finally, we processed the EV-depleted plasma, collected after EV isolation, for cfDNA extraction as previously described (**Fig. 1a** and **Fig. 2a**). The concentration of cfDNA was comparable in the two BrCa subtypes (mean HER2+: 3.9ng/mL; mean HER2-:2.9 ng/mL) within the early-stage cohort (**Fig. 2f** and **Supplementary Fig. 3e**).

Furthermore, in the same BrCa patient cohort, no correlation was observed between the amounts of cfDNA and EVs, regardless of the HER2 status obtained on tissue biopsies according to clinical classification (**Fig. 2g**).

Altogether our data suggested that HER2+ and HER2- patients exhibit comparable quantity and quality of EVs and cfDNA, thereby supporting the opportunity of profiling both analytes for cancer-associated biomarkers.

### cfDNA and EVs represent circulating components that reflect the patient tumor molecular features assessed on tissue biopsies

We sought to provide proof of principle validation of ONCE–derived analytes for cancer cargo detection in liquid biopsies by taking advantage of the possibility of jointly screening cfDNA with EV cargo. To detect molecular alterations associated with BrCa, we performed DNA whole-exome sequencing (WES) (mean coverage = 537x, min=282x, max=731x) and RNA deep sequencing (mean depth=88M reads, min 56M reads, max=97M reads, detecting 13009 transcripts on average at cpm>5, min=8545, max=19113) respectively on cfDNA and EV-RNA isolated from two HER2+ (PMB2.8 and PMB2.36) and two HER2- (PMB2.26 and PMB2.30) BrCa patients (**Supplementary Table 1** for patient marker status, **Supplementary Table 2** for WES coverage, **Supplementary Table 3** for cfDNA and EVs quantification and **Supplementary Table 4** for RNAseq statistics).

As illustrated by the circos plots (**Fig. 3a**), DNA-Seq of the cfDNA samples detected several cancer-related genomic alterations and confirmed the amplification of *ERBB2* (on chromosome 17) of patients PMB 2.8 and PMB 2.36, in line with diagnostic tissue biopsy *in situ* assessment (**Supplementary Table 1**). Multiple cancer–related genomic alterations, but no *ERBB2* amplification, were present in the cfDNA samples from patients PMB2.30 and PMB2.26, classified as HER2- (**Fig. 3a**). RNA-Seq of the EV-RNA from the same patients showed that transcripts encoding for HER2 were detectable on samples PMB2.8 and PMB2.36, both HER2+ (**Fig. 3b**), but not on the sample PMB2.30 (HER2-). Interestingly, the sample PMB2.26 was classified as HER2- based on the absence of *ERBB2* genomic amplification by FISH assay. However, the corresponding IHC on the tissue biopsy showed low to moderate membrane immunoreactivity to HER2 antibody in 50% of tumor cells (**Supplementary Table 1**), compatible with the EV-RNA sequencing data, and overall suggesting a potential HER2 overexpression regulation at the transcriptional or post-transcriptional level [31–35]. Additionally, transcripts encoding for markers of prognostic value in BrCa, such as MKI67 (a marker of proliferation Ki-67) and estrogen receptor alpha (ESR1) [36], were differentially detected in both HER2+ (PMB2.8 and PMB2.36) and HER2- samples (PMB2.30 and PMB2.26) (**Fig. 3b**; Plasma) in line with tissue biopsy data (**Fig. 3b**; Tissue /Protein). Altogether, the sequencing data, despite the limited sample size, confirmed cfDNA obtained through ONCE as a circulating source of information for cancer profiling and highlighted the relevance of the EV-RNA cargo as a powerful additional component for cancer biomarkers detection in multi-analyte liquid biopsies. Additionally, by western blotting analysis on EV-associated proteins released in the medium by HER2+ (SKBR3) and HER2- (MDAMB231) breast cancer cell lines, we assessed EVs as carriers of the HER2 biomarker in the medium from HER2+ cancer cell line only (**Supplementary Fig.4a**). By using immuno-mediated staining in combination with imaging flow cytometry, we were able to confirm the association of HER2 to the surface of EVs subpopulations and their differential abundance among a subset of n=3 BrCa HER2+ and n=3 HER2- samples (**Fig. 3c and Supplementary Fig. 4b for IgG isotype control**). In conclusion, in this proof-of-principle experiments, we assessed that ONCE-derived cfDNA and EVs represent an informative source for investigating cancer biomarkers, such as HER2, in the circulation, as genomic and molecular features are representative of matched tissue-based investigations.

**Figure 3:**
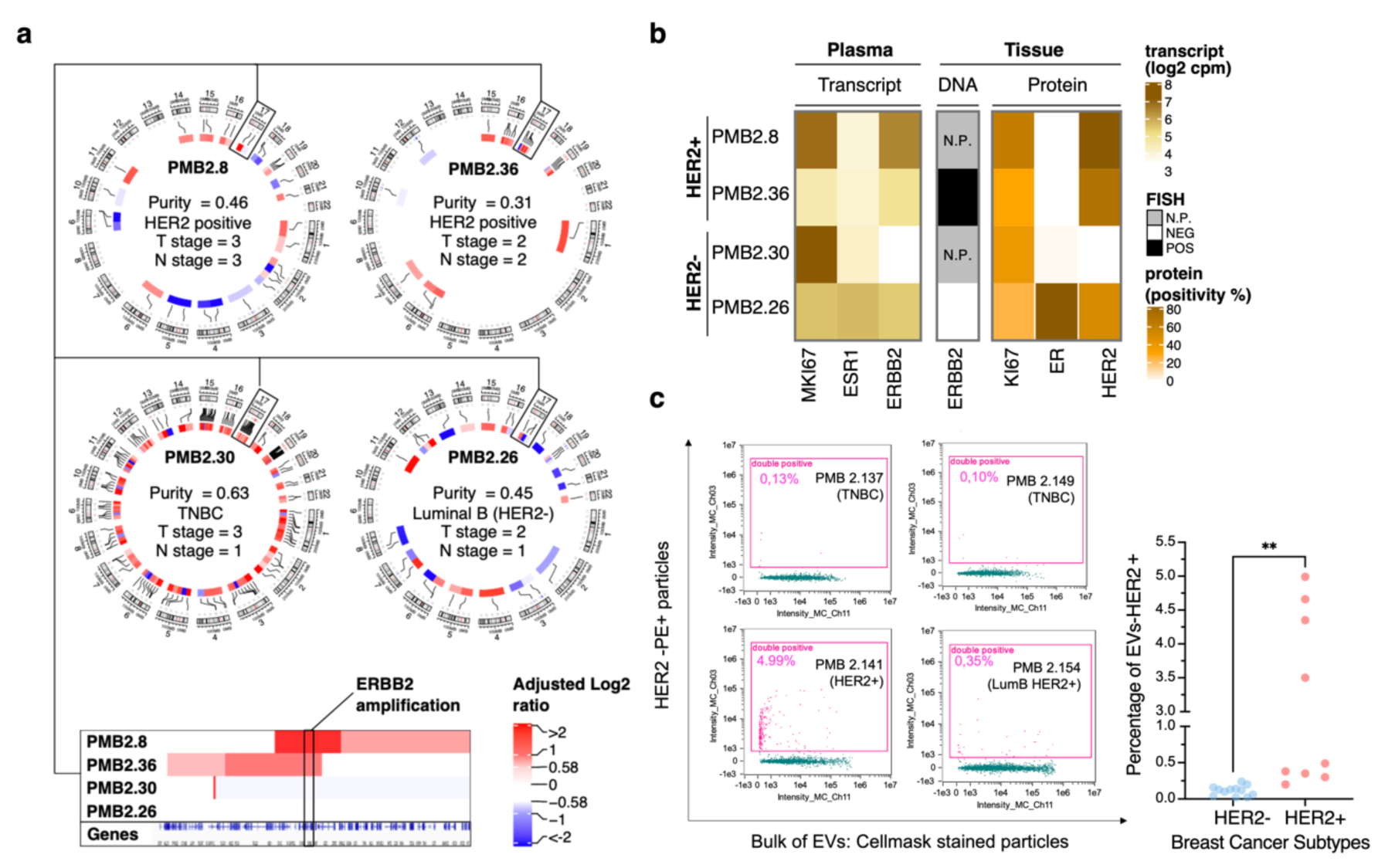
Circulating cfDNA and EVs in early stage BrCa patients are a remarkable source for detecting genetic and molecular alterations. **a.** Circos plots of cfDNA analyzed by whole-exome DNA-Seq (WES) of n=4 representative BrCa patients. Squares outline the *ERBB2* gene locus (Chr.17q12) shown as linear visualization in the inset; *ERBB2* gene amplification is detectable on samples PMB2.8 and PMB2.36, (cfDNA isolated from HER2+ BrCa patients); no *ERBB2* amplification is detectable in cfDNA samples PMB 2.30 and PMB2.26 (cfDNA isolated from HER2- BrCa patients). Purity indicates the circulating tumor content. **b.** Heatmap showing the detection of relevant breast cancer biomarkers in n=4 patients: PMB2.8 and PMB2.36, (HER2+), PMB2.30 and PMB2.26 (HER2-). Transcripts were detected by RNA-Seq from EV-RNA from liquid biopsies. Proteins were detected by IHC on tissue biopsies. DNA amplification of *ERBB2* gene was detected by FISH on tissue biopsies. *ERBB2* (HER2), ESR1 (Estrogen Receptor alpha) and MKI67 (Marker of Proliferation Ki-67). **c.** Representative plots and quantification of HER2+ EVs subpopulations by imaging flow cytometry (Amnis Imagestream^x^ MK II). Plots show CellMask staining utilized to visualize the bulk of lipid nanoparticles on x-axis (Intensity_MC_Ch_11) and PE-HER2 Antibody staining on y-axis (Intensity_MC_Ch_03). Data are from technical replicates of measurements performed on EVs samples isolated from n=3 HER2- and n=3 HER2+ BrCa patients (PMB). ** p=0.0041 by t-test. **cfDN**A: cell free DNA; **EVs**: Extracellular Vesicles; **EV-RNA**: RNA extracted from EVs; **BrCa**: Breast Cancer; **HER2+**: HER2 positive, **HER2-**: HER2 negative; **IHC**: Immunohistochemistry. See also Supplementary Fig. 4

### Integrating EV-RNA and cfDNA data increases precision in identifying the HER2 status at an early stage of BrCa

After demonstrating that cfDNA and EVs can be isolated from BrCa patients with no significant differences in the recovered amount across the two subtypes **(Fig. 2b and 2e,f**) and that they allow the detection of cancer–associated alterations (**Fig. 3a,b**), we sought to quantify the ability of such circulating nucleic acids to screen for the status of a critical BrCa biomarker, such as HER2 in early stage BrCa liquid biopsies. We hypothesized that the combination of both EV-RNA and cfDNA-based quantitative assessments would increase the detection performance by overcoming potential limitations on a patient basis (e.g., a low fraction of circulating tumor DNA, a low proportion of cancer-related EVs in the circulation).

We analyzed EV-RNA and cfDNA by ddPCR, a fast technique widely applied to screen large numbers of samples and that has already been successfully used in clinical liquid biopsies-based analyses for its high sensitivity and specificity [37, 38]. To ensure a large enough set of BrCa patient plasmas with EV-RNA and cfDNA yields adequate to run the two ddPCR assays, we extended the study cohort to additional patients from the same prospectively collected clinical cohort (**Table 1**). Thus, we characterized a set of n= 38 BrCa (HER2+ and HER2-) samples for *ERBB2* gene amplification on cfDNA and *ERBB2* transcript expression on EV-RNA. Amplification on cfDNA was determined as the ratio between circulating *ERBB2* and reference gene *EIF2C1* by using a previously reported ddPCR assay for *ERBB2* copy number alteration on BrCa tissues [39]. The expression on EV-RNA was calculated as a ratio between the detection of fragments corresponding to ERBB2 and the commonly utilized tissue reference gene *EEF2* [40]. To evaluate the specificity of this ddPCR assay, we included a set of n=7 HDs (to define the thresholds on *ERBB2* amplification and overexpression) and EV-RNA extracted from conditioned cultured medium of HER2+ (SKBR3) and HER2- (MDAMB231) breast cancer cell lines. ddPCR data showed that 57% of the HER2- BrCa patients (**Fig.4a**, blue dots) group with the HDs, while 83% of HER2+ BrCa patients (**Fig.4a**, red dots) show higher signals than HDs on *ERBB2* on EV-RNA (*ERBB2/EEF2*; **Fig. 4a, y-axis**) and cfDNA (*ERBB2/EIF2C1;* **Fig.4a, x-axis**). HER2- patients never demonstrate cfDNA *ERBB2* gene amplification, whereas 8 of them show mild transcript overexpression in the lowest range of expression of HER2+ patients. ddPCR data on EV-RNA and cfDNA showed concordant marked positivity for about 30% of the HER2+ BrCa samples. In comparison, 25% (n= 6 out of 24 red dots) and about 20% (n= 5 out of 24 red dots) of HER2+ samples were correctly classified by EV-RNA only and cfDNA only, respectively. EV-RNA resulted of particular importance to capture samples that were classified positive for HER2 based on *ERBB2* gene amplification by FISH on tissue biopsies (**Fig. 4b**), but for which the amplification was not detectable in cfDNA (as PMB2.19, PMB2.70, PMB2.89).

**Figure 4:**
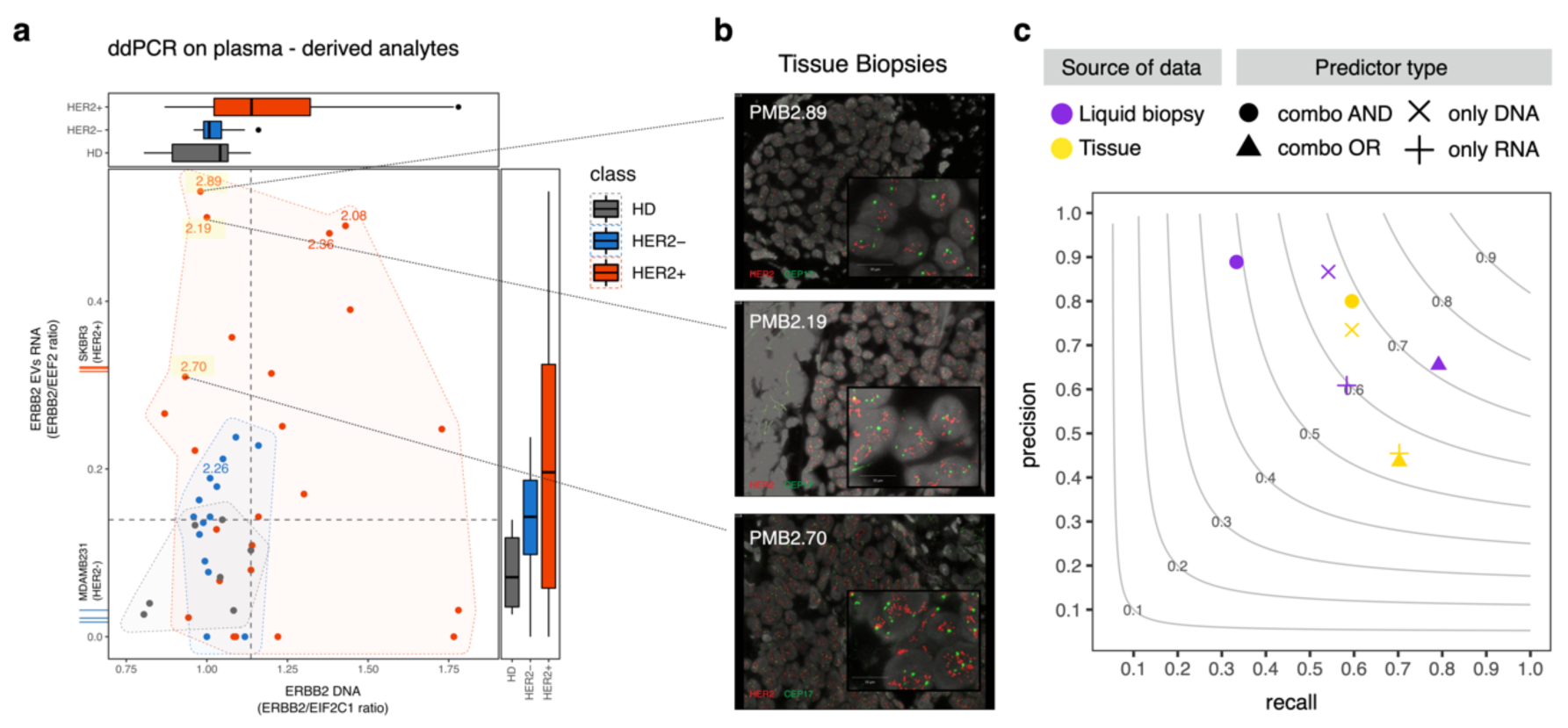
Integrating data from circulating cfDNA and EV-RNA strengthens the potential of liquid biopsies in early stage BrCa. **a.** Concomitant ddPCR of *ERBB2* gene on cfDNA and transcript on EV-RNA. Red dots: HER2+ BrCa (n=24); blue dots: HER2- BrCa (n=14) (stratification based on tissue biopsies data); grey dots: HDs (n=7). Marginal boxplots show distributions for each group of samples. Labeled samples are shown in panel b. Colored areas show the area that contains all the points of a specific class of samples (HDs, HER2+ or HER2-). Grey lines mark thresholds based on HDs distributions for HER2 positivity prediction (maximum level of RNA and DNA detected in HDs is set as threshold). Measurements from EV-RNA of SKBR3 (HER2+) and MDAMB231 (HER2-) breast cancer cell lines (n=3 replicates) are indicated on the Y axis by red and blue lines, respectively. **b.** Fluorescent In Situ Hybridization (FISH) pictures of breast tumor tissue biopsies show HER2 gene amplification. Quantification of amplification was based on HER2 (red) and CEP17 (green) ratio according with the American Society of Clinical Oncology (ASCO)/College of American Pathologists (CAP) guidelines. Data are associated to BrCa patients: PMB2.89 (HER2/CEP17 ratio: 3.0), PMB2.70 (HER2/CEP17 ratio: 5.8), PMB2.19 (HER2/CEP17 ratio: 2.9). **c.** Performance of HER2 status classifiers: combining cfDNA and EV-RNA data increases the performance of liquid biopsy biomarker detection. Liquid biopsy classifier (violet) is obtained using data from panel a. Classifier for Tissue biopsies (yellow) uses tissue data from Supplementary Fig. 5. Symbol shape indicates if the classifier uses only DNA or RNA data or a combination of both. Samples with values higher than the thresholds are classified as HER2+ based on one of the following rules: 1. only RNA: only the signal of RNA reaches the threshold; 2. only DNA: only the signal of DNA reaches the threshold; 3. Combo AND: both DNA and RNA signals reach the threshold; 4. Combo OR: signal for any DNA or RNA reaches the threshold. Precision is the ratio between predicted true positives and the total number of predicted positives. The recall is the ratio between predicted true positives and the total number of positives in the population. Lines in the background show the F1 score (harmonic mean between recall and precision). The F1 score is a score of measurements’ performance and ranges from 0 (lowest recall; lowest precision) to 1 (highest recall; highest precision). **cfDN**A: cell-free DNA; **EVs**: Extracellular Vesicles; **EV-RNA**: RNA extracted from EVs. **ddPCR**: droplet digital PCR; **HDs**: Healthy Donors, **BrCa**: Breast Cancer; **HER2+**: HER2 positive, **HER2**-: HER2 negative; **HDs**: healthy donors. See also Supplementary Fig. 5.

To estimate the relevance of combining EV-RNA and cfDNA signal in BrCa, we compared the classification performance of every single analyte (“*only DNA*”, “*only RNA*”) versus combined analytes (“*combo OR*”, “*combo AND*”, i.e., high level of either one or both analytes is required for positivity; see **Methods**) by focusing on the presence or absence of *ERBB2* amplification or HER2 overexpression (see method section **HER2 positivity classification of plasma and tissue samples**). The prediction accuracy was measured by the F1 score (**Fig. 4c**, grey lines), that is the harmonic mean between the recall (the fraction of HER2+ cases that can be detected; **Fig. 4c, x-axis**) and the precision (the fraction of cases that are correctly classified as HER2+; **Fig. 4c, y-axis**); the higher the F1 score, the better the classification accuracy. To compare liquid biopsy-based results and tissue-based results, we similarly quantified the classification performance from DNA and/or RNA sequencing data from the TCGA BrCa patient tissue cohort (n=537) [41].

We observed that the highest performance is obtained by the combination of the two analytes (*combo OR*, violet triangle) and that the results obtained from the liquid biopsy are not inferior to those obtained from tissue (yellow triangle) (**Supplementary Table 5**). Thus, using the information of both analytes in liquid biopsy (*combo OR*, violet triangle) leads to classification performance comparable to the *combo AND* approach in tissues (yellow circle), which is the best-performing predictor for tissue data.

Altogether these data demonstrate that the power of ONCE–derived EV-RNA and cfDNA analysis for capturing HER2 positive cases is comparable to that obtained by sequencing of tissue nucleic acids.

## Discussion

Integrating multiple circulating analytes information is challenging [42]. The proposed ONCE protocol allows the implementation of liquid biopsies-based tests limiting the blood volume to be collected from patients and, at the same time, improving operational and analytical procedures. This is particularly relevant for pediatric and severely debilitated patients, for which accessibility to blood draws may be limited. Processing single aliquots of plasma is less time-consuming and more cost-effective as it halves the cost of blood draws and subsequent laboratory processing compared to using multiple aliquots. Furthermore, on the analytical side, using a single aliquot to analyze two components allows discriminating among technical and biological effects, such as the possibility to distinguish if low levels of circulating cfDNA or EVs amounts are likely due to degradation issues versus low shedding rate of the tumor cells. Notably, the high quality of ONCE-isolated circulating nucleic acids (cfDNA and EV-RNA) is suitable for ultra-sensitive techniques such as the ddPCR and NGS and the subsequent potential screening of genetic alterations at early-stage of cancer.

By proposing the analysis of EVs as a circulating component in combination with cfDNA, the ONCE protocol enables the detection of cancer biomarkers in patients where the relevant biomarker signal presents at the transcriptional level and would be missed by analysis of cfDNA only. Indeed, we showed that the ONCE protocol might be an informative approach to monitor *ERBB2* in breast cancer patients without genomic *ERBB2* amplification or patients that are HER2 positive at diagnosis and turn negative after multiple cycles of therapy [43, 44]. Furthermore, the combination of two diverse analytes, the cfDNA and EV–RNA in the context of non-metastatic early-stage BrCa patients represents an effective strategy for enabling liquid biopsies when the circulating tumor content is limited and restrains the sensitivity in detecting biomarkers and circulating oncogenic transcriptional variants [45].

In addition to the EV-RNA component, the advantage of including EVs as a player in multi-analyte liquid biopsy approaches is represented by the variety of molecular features embedded into their cargo that allows extending cancer biomarkers analysis to other biomolecules, such as proteins exposed on their surface. Using imaging flow cytometry, we identified subpopulations of EVs carrying HER2 and estimated their relative abundance compared to the total number of circulating EVs. A further implementation of our approach by combining multiple cancer biomarkers and/or platforms to screen large numbers of samples in short time (i.e., ELISA assays) would represent a significant advance for investigating EVs in liquid biopsies.

Additionally, diverse methods for EV isolation can be adapted to the ONCE protocol and selected based on recommendations proposed by MISEV 2018 [46] and by specific research interests on EVs as companion analytes for liquid biopsies.

Altogether the data here presented confirm the possibility of assessing the status of multi-analyte biomarkers in early-stage disease from a single aliquot of plasma by investigating the circulating EVs and cfDNA.

Further studies will be essential for validation on more extensive sample collections and diverse tumor types for the future implementation of multi-analyte liquid biopsy-based assays in the clinical setting.

## Author Contributions

VM, YC, VDA, CN, and FD conceived the study, analyzed the results, and wrote the manuscript. VM, and YC performed the experiments and generated the data. OQ, ST, MN, FV, AM, EL, SP, AM, DR, and IP contributed to the methodology, technology, and experiments. SSM, MB, AF, OC, GA, and DDV contributed to the investigation and the manuscript editing. All authors interpreted the data, and approved the final version.

## Acknowledgements

We thank all members of the Demichelis laboratory for critical comments. We are grateful to the High Throughput Screening (HTS), Cell Analysis and Separation (CASC) and Next Generation Sequencing (NGS) core facilities at CIBIO and to Fabio Malesani, Salvatore Girlando, and Cristina Del Bianco for technical support with the experiments. We are also thankful to Roberta Tarallo for helping with the processing of the blood samples.

## Data sharing statement

The datasets used and/or analyzed during the current study are available from the corresponding author upon request.

## Supplementary Figures

**Supplementary Figure 1 related to Main Figure 1.**
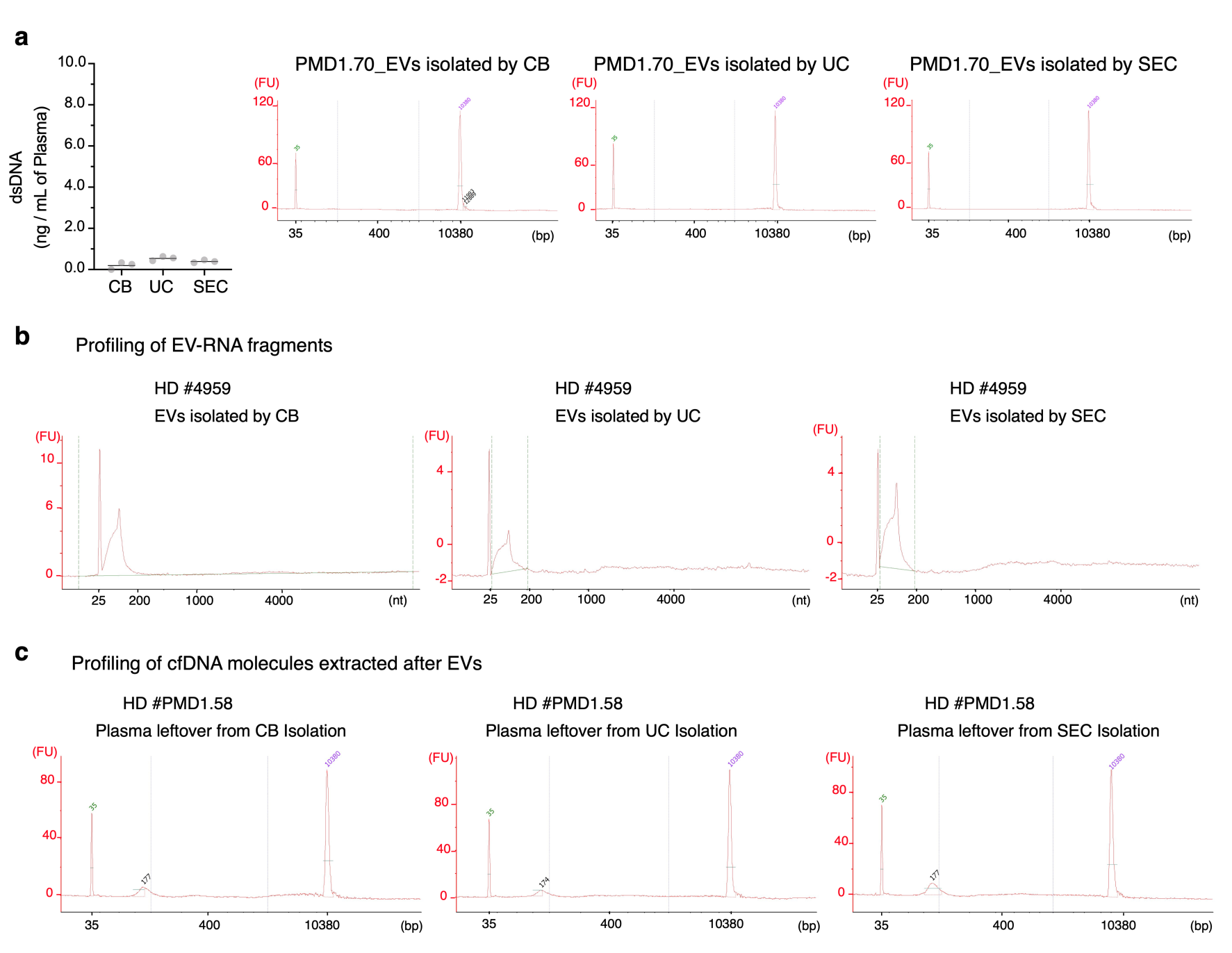
Short fragments of nucleic acids are isolated from plasma. **a.** Quantification of EV-associated dsDNA and representative bioanalyzer profiles obtained by Agilent HS DNA bioanalyzer assay. The dsDNA was quantitated by Qubit dsDNA HS assay after extraction from EV samples isolated by CB, UC or SEC. EV samples were from plasma (1.8mL) of n=3 HDs. Bioanalyzer plots were from healthy donor PMD1.70. **b.** Representative profiles of RNA samples isolated from EVs (EV-RNA) obtained by Agilent RNA 6000 Pico bioanalyzer assay. Bioanalyzer internal standard is at 25bp. EVs were separated from plasma of HD #4959 by using three diverse methods: CB, UC or SEC. **c.** Representative bioanalyzer profiles of cfDNA obtained by Agilent HS DNA bioanalyzer assay. cfDNA was extracted from plasma leftovers collected after EVs isolation by CB, UC or SEC. Internal standards are at 35bp and 10380bp. Plasma aliquots (1.8mL) were from HD #PMD1.58. EV: Extracellular Vesicles; CB: Charge – Based EV Isolation Method; UC: Ultracentrifugation; SEC: Size Exclusion Chromatography; dsDNA: double-stranded DNA; EV-RNA: RNA samples extracted from EVs; cfDNA: cell free DNA; HDs: Healthy Donors.

**Supplementary Figure 2 related to Main Figure 1.**
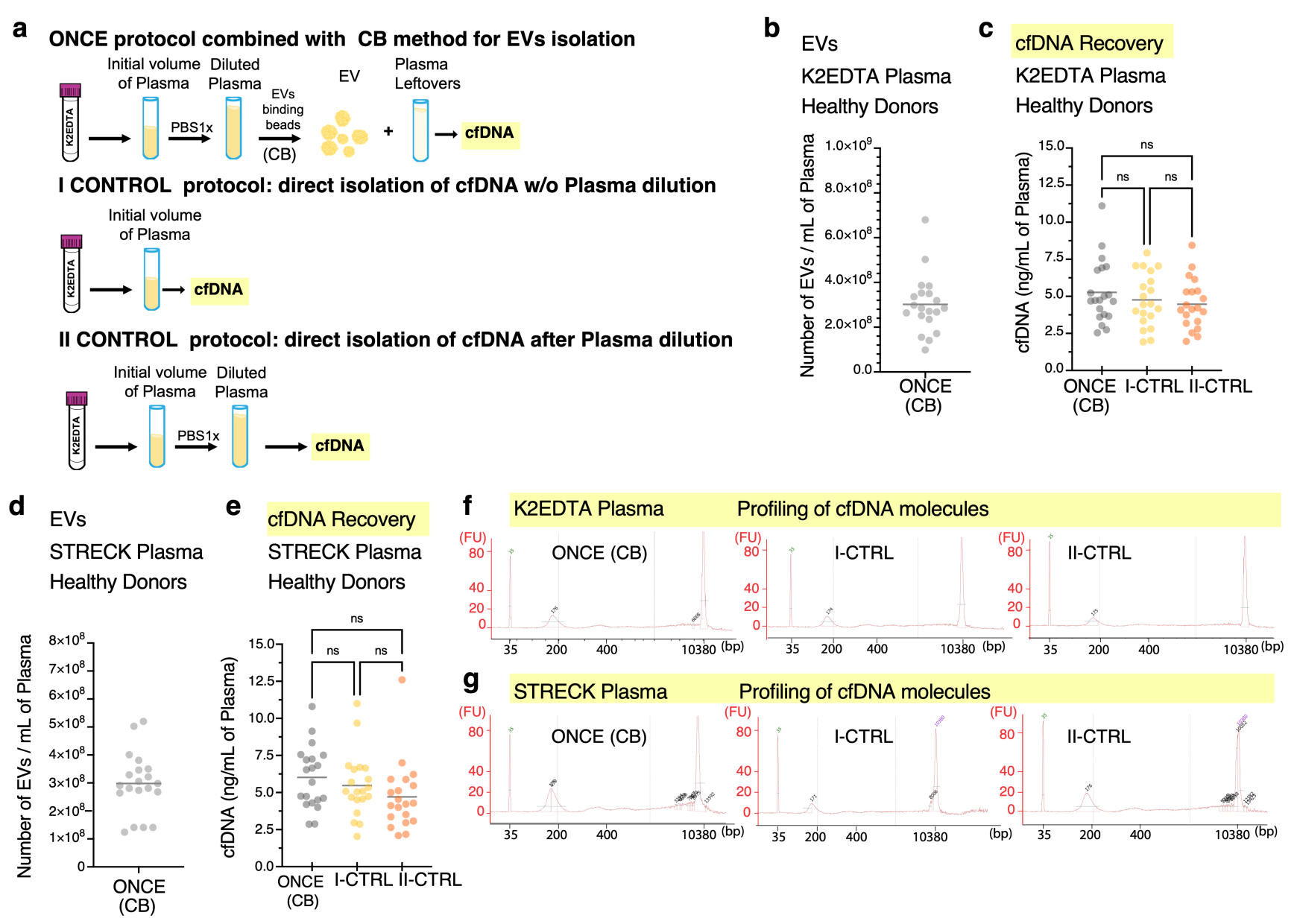
ONCE protocol, combined with CB, allows efficient isolation of EVs and cfDNA from plasma of blood collected into EDTA or Streck tubes. **a.** Scheme of ONCE protocol performed by applying the CB method for EVs isolation. ONCE protocol is the sequential isolation of EVs and cfDNA from the same aliquot of diluted plasma. I CONTROL (I-CTRL) is the original method for cfDNA isolation from a single aliquot of not–diluted plasma. II CONTROL (II – CTRL) is the isolation of the cfDNA from a single aliquot of diluted plasma. Plasma dilution is an essential step to reduce plasma viscosity thereby facilitating the binding of the beads to the EVs. **b.** Quantification of EVs by TRPS measurements. EVs were isolated by CB from plasma of blood collected in K2EDTA tubes. EVs were from a cohort of n=20 HDs. **c.** Quantification of cfDNA obtained by Qubit dsDNA HS assay (Thermo Fisher Scientific). cfDNA samples were extracted from plasma leftover collected after EV isolation. Data are from n=20 HDs. An aliquot of blood was collected from each individual donor in K2EDTA tubes and processed for plasma separation. The volume of plasma was split into three aliquots. Each aliquot was processed with a dedicated protocol (either one of ONCE, I-CTRL or II-CTRL II) as shown in panel a. n.s.: not significant differences by One – way Anova. **d.** Quantification of EVs by TRPS measurements. EVs were isolated by CB from plasma of blood collected in Streck tubes. EVs were from a cohort of n=20 HDs. **e.** Quantification of cfDNA by Qubit dsDNA HS assay (Thermo Fisher Scientific). cfDNA was extracted from plasma leftovers collected after EV isolation. Samples are from a cohort of n=20 HDs. An aliquot of blood was collected from each individual donor in Streck tubes and processed for plasma separation. The volume of plasma was split into three aliquots and each of them was processed with a dedicated protocol (either one of ONCE, I-CTRL or II- CTRL II) as shown in panel a. n.s.: not significant differences by One – way Anova. **f.** Representative profiles of cfDNA obtained by Agilent HS DNA bioanalyzer assay. cfDNA samples were extracted from plasma of blood collected into K2EDTA tubes. Samples were processed according to protocols ONCE, I- CTRL, II- CTRL as shown in panel A. All profiles are comparable demonstrating that the length of fragments of cfDNA isolated from diluted plasma after EVs recovery (ONCE) is comparable to cfDNA obtained from not diluted plasma (I- CTRL) or from diluted plasma without processing for EVs isolation (II-CTRL). The typical cfDNA fragment size (about 174-176 bp) is evident across all profiles. Internal standards are at 35bp and 10380bp. **g.** Representative profiles of cfDNA obtained by Agilent HSDNA bioanalyzer assay. cfDNA samples were extracted from plasma of blood collected into Streck tubes. Samples were processed according to protocols ONCE, I- CTRL, II- CTRL as shown in panel A. All profiles are comparable demonstrating that the length of fragments of cfDNA isolated from diluted plasma after EVs recovery (ONCE) is comparable to cfDNA obtained from not diluted plasma (I- CTRL) or from diluted plasma without processing for EVs isolation (II-CTRL). The typical cfDNA fragment size (about 174-176 bp) is evident across all profiles. Internal standards are at 35bp and 10380bp. ONCE: ONe Aliquot for Circulating Elements; EV: Extracellular Vesicles; cfDNA: cell free DNA; HDs: Healthy Donors; TRPS: Tunable Resistive Pulse Sensing; CB: Charge – Based EV Isolation Method; I- CTRL: First Control Protocol; II-CTRL: Second Control Protocol; cfDNA: cell free DNA, K2EDTA tube: tube containing di-potassium K2EDTA which blocks the coagulation cascade. K2EDTA tubes are commonly used for examination of whole blood in haematology. STRECK tube: Cell-Free DNA BCT STRECK is a blood collection tube that which stabilizes nucleated blood cells preventing release of genomic DNA and allowing isolation of high-quality cell-free DNA. Cell-Free DNA BCT tubes (cfDNA BCTs) are commercialized by Streck (La Vista, NE).

**Supplementary Figure 3 related to Main Figure 2.**
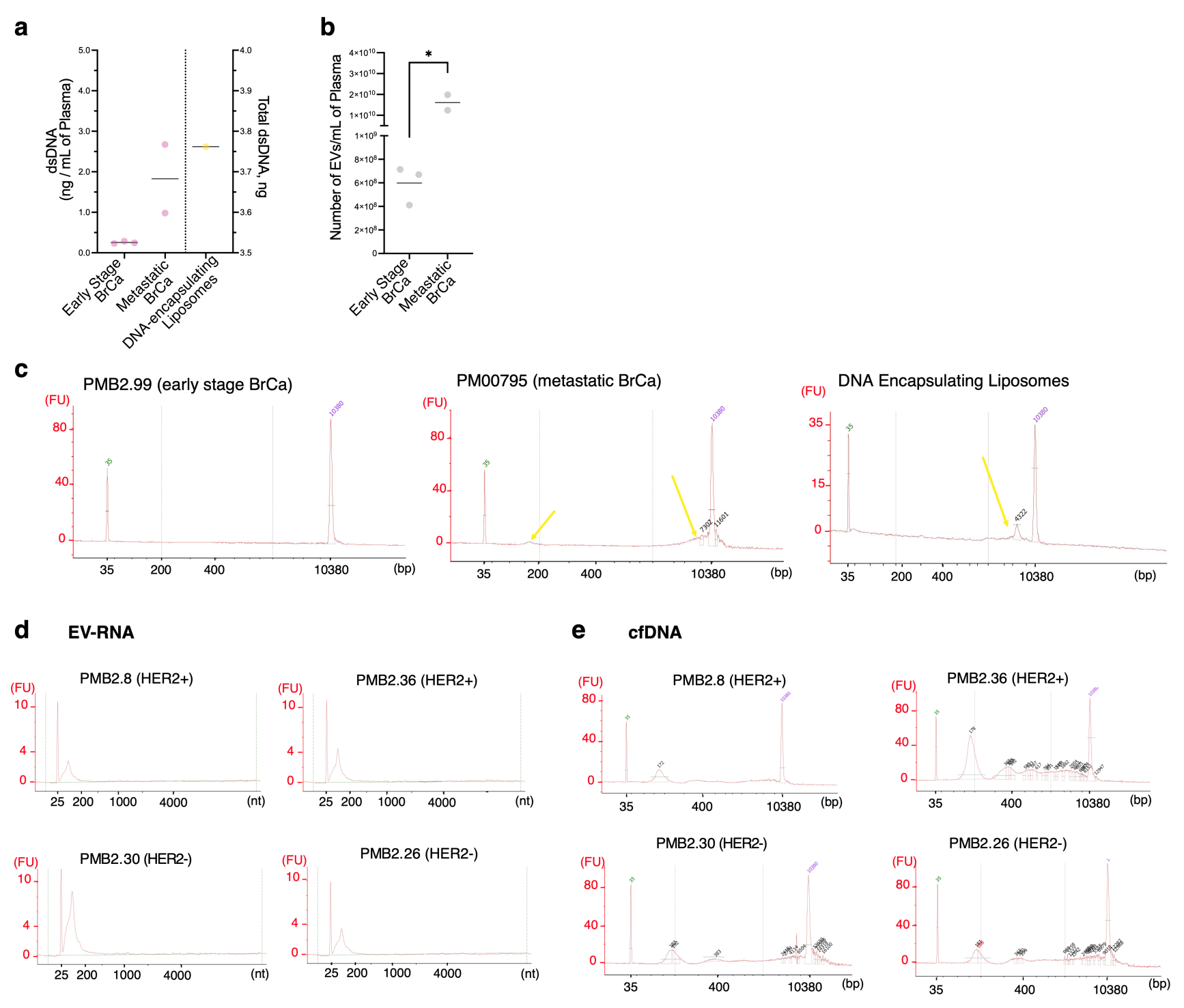
Small EVs isolated from BrCa patients at early stage contain minuscule amount of dsDNA. **a**. Quantification of EV-associated dsDNA by Qubit dsDNA HS assay. dsDNA was extracted from EVs after isolation by CB. EVs were from early stage or metastatic BrCa patients. Synthetic liposomes (100-200nm diameter size) containing artificial dsDNA (DNA-encapsulating liposomes) were utilized as internal control. The concentration of dsDNA recovered from early stage BrCa patients is miniscule, not suitable for ddPCR and/or sequencing assays. **b.** Quantification of EV samples utilized for dsDNA extraction. Measurements were performed by NTA. EVs were isolated from plasma aliquots of BrCa patients at early stage (n=3) or advanced, metastatic stage (n=2). The concentration of EVs is significantly higher on samples derived from metastatic patients. * p value= 0.0112 by unpaired t-test. **c.** Representative EV-DNA profiles obtained by Agilent HS DNA bioanalyzer assay. Samples are EVs from plasma of metastatic or early stage BrCa patients or synthetic liposomes containing artificial dsDNA (internal control). Yellow arrows indicate signals associated to DNA. **d.** Profiling of sequenced EV-RNA samples by Agilent RNA 6000 Pico bioanalyzer assay. Internal standard is at 25bp. Labels report BrCa patient ID and subtype. All EV-RNA were extracted from EVs isolated by CB from plasma of early stage BrCa (PMB). **e.** Profiling of sequenced cfDNA samples by Agilent HS DNA bioanalyzer assay. Labels report BrCa patient ID and subtype. All cfDNA were isolated in the framework of ONCE from plasma leftover of early stage BrCa (PMB). Internal standards are at 35bp and 10380bp. EV: Extracellular Vesicles; EV-DNA: DNA extracted from EVs; EV-RNA: RNA extracted from EVs; dsDNA: double-stranded DNA; BrCa: Breast Cancer; NTA: Nanoparticle Tracking Analysis.

**Supplementary Figure 4 related to Main Figure 3.**
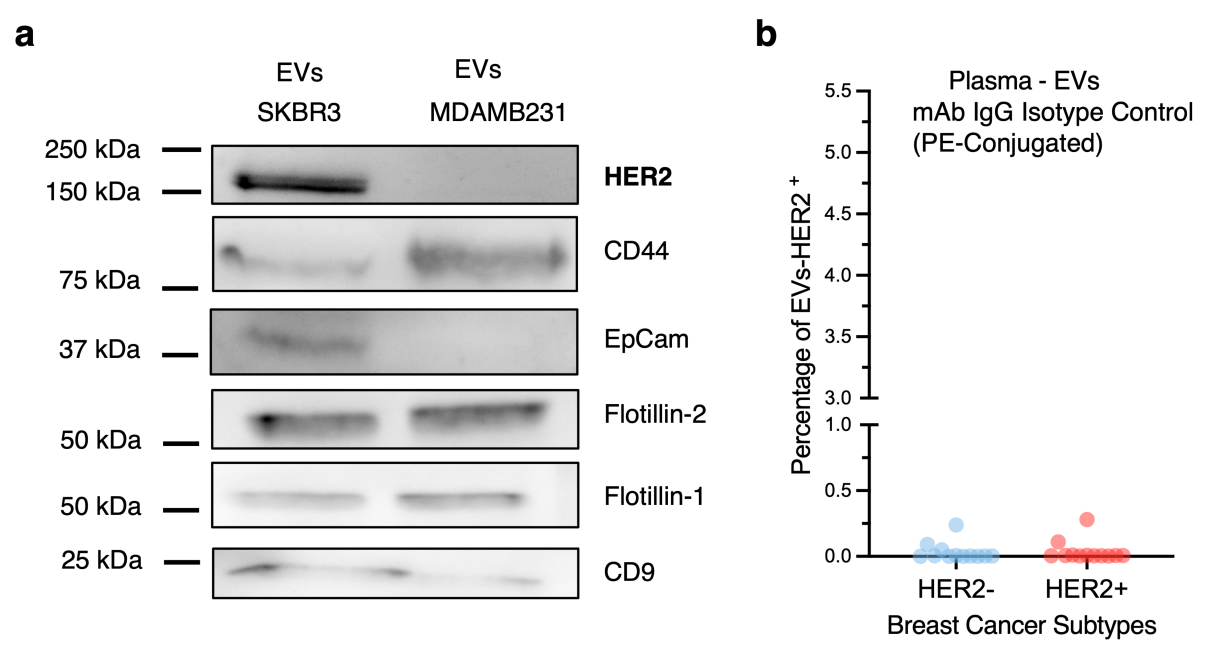
HER2 protein is carried by EVs. **a.** Western Blot assay showing the breast tissue associated proteins HER2, CD44, EpCaM and EV-enriched proteins Flotillin 1, Flotillin 2 and CD9 in representative n=2 protein extracts of EV samples isolated by CB from conditioned medium of SKBR3 (HER2+) and MDAMB231 (HER2-) human breast cancer cell lines. **b.** Quantification of PE+ signal detected by imaging flow cytometry (Amnis Imagestreamx MK II). EV samples were stained with the IgG isotype control (PE-Conjugated) for the anti-HER2 antibody utilized in Fig. 3c. As expected, only minimal and comparable percentages of fluorescent particles are detected on both HER2- and HER2+ samples. Data are from technical replicates of measurements performed on EVs samples isolated from n=3 HER2- and n=3 HER2+ BrCa patients.

**Supplementary Figure 5 related to Main Figure 4.**
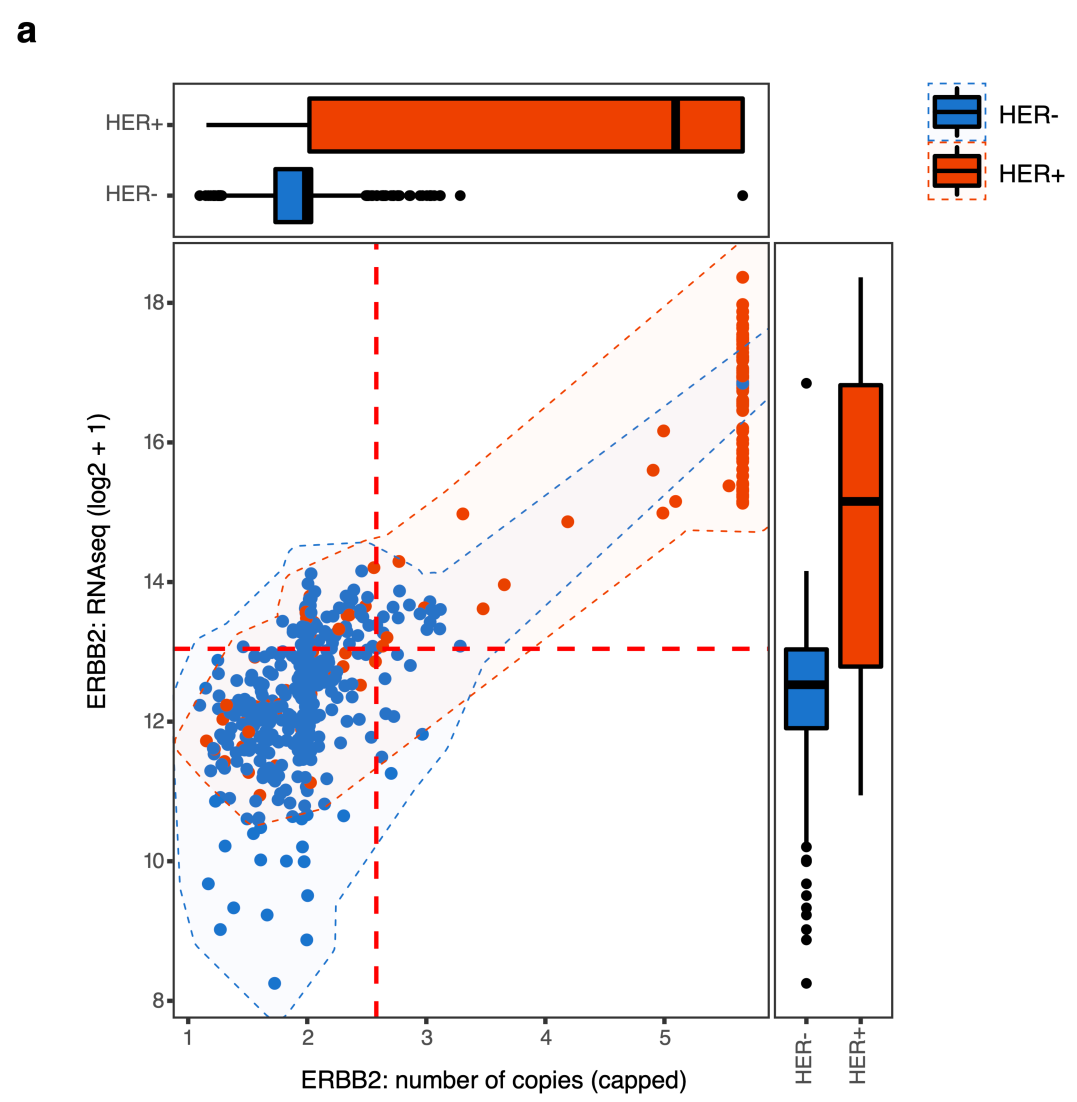
Tissue based ERBB2 status of the TCGA BrCa cohort. **a.** Plot derived from tissue based sequencing data of BrCa patients deposited on the cancer genome atlas (TCGA) data portal. X axis: ERBB2 Number of copies by DNA-Seq; Y axis: expression of ERBB2 gene as quantified by RNA-Seq datasets/studies (Ciriello G, Gatza ML, Beck AH, Wilkerson MD, Rhie SK, Pastore A, Zhang H, et al. Comprehensive Molecular Portraits of Invasive Lobular Breast Cancer. Cell 2015;163:506-19). Specifically, data from RNA-Sequencing and DNA Sequencing were analyzed to evaluate how DNA-associated or RNA – associated information predict the HER2 Status (positive vs negative). Red and blue colors show stratification of BrCa by IHC tissue biopsy (protein – associated data). Marginal boxplots show distributions for each group of samples. Horizontal and vertical dotted lines indicate thresholds to define ERBB2 gain or over-expression.

## Supplementary Methods

### Experimental design and human sample collection

Healthy donors (n=20; males and females, age >18 years old with no medical history of cancer or anti-cancer treatments) prospectively enrolled on a voluntary basis on a protocol approved by the University of Trento Ethics Committee (ID # 2017-010). Healthy donor plasma samples were used to test diverse EV isolation methods, followed by cfDNA extraction within the ONCE framework.

Breast cancer patients’ plasma samples were obtained from a cohort of patients prospectively enrolled on a protocol approved by the Ethics Committee of Santa Chiara Hospital in Trento (Rep.Int.12315 of July 24, 2017) with written informed consent. Eligibility criteria included breast cancer diagnosis and recommendations for neoadjuvant therapies. A set of n=44 patients to similarly represent each of the four most common subtypes of breast cancer ((HER2+, Luminal B (HER2+), Luminal B (HER2-), TNBC)) was initially selected to compare the circulating tumor–derived content (EVs and cfDNA) from HER2 positive and HER2 negative breast cancer samples at diagnosis. To then ensure a large enough set of patients with adequate EV-RNA and cfDNA yields for the ddPCR assays, additional patients were selected from the same prospectively collected clinical cohort (ddPCR successfully performed on n=38 BrCa patient plasmas), and n=7 commercially available healthy donor plasma samples (males and females, age > 40 years old with no medical history of cancer or anti-cancer treatments) were purchased from Precision Medicine Group, LLC as controls.

### Blood processing and plasma isolation

Whole peripheral blood (8.0 –10 ml) was collected into K2EDTA-containing tubes (Vacuette, Greiner Bio-One) or Streck tubes (Streck Cell-Free DNA BCT; Streck; La Vista, NE, USA) for comparison. Plasma from K2EDTA tubes was isolated within 2 hours from the collection by two-step centrifugation at room temperature (1700 rcf x 15 min; 3000 rcf x 10 min). Plasma from Streck tubes was isolated within five days from the collection by two-step centrifugation at room temperature (1700 rcf x 15min; 15000 rcf x 20 min). After isolation, plasma was immediately processed or aliquoted and stored at −80°C.

### ONCE Protocol: combined isolation of EVs and cfDNA from human plasma

#### EV isolation by Charge-Based (CB) method and cfDNA extraction

The human plasma samples were filtered (Minisart NML syringe filters, pore size: 0.8 µm; Sartorius) and diluted with 1x Phosphate-Buffered Saline without calcium and magnesium buffer (Gibco) at 1:3 (v/v) ratio and processed for EV isolation by a charge – based isolation method as described before [1, 2]. The diluted plasma was recovered and processed for cfDNA isolation by QIAmp Circulating Nucleic Acid kit (Qiagen) [3].

To verify the efficiency of cfDNA extraction after EV isolation by CB method, we collected blood from n=20 HDs. Two types of tubes commonly utilized for liquid biopsies, i.e., K2EDTA and BCT Streck tubes, were used. Each volume of the separated plasma sample was split into three aliquots (1.5ml/each). The three aliquots of plasma were then processed, each with one of the three following protocols as described in **Supplementary Fig. 2a**:

i) ONCE Protocol, the plasma aliquot was diluted in PBS1X and incubated with EV-capture beads for EV isolation; the diluted EV-depleted plasma leftovers are then utilized for cfDNA extraction.
ii) I CONTROL protocol, the aliquot was processed to purify cfDNA according to the protocol commonly utilized for cfDNA isolation [4];
iii) II CONTROL protocol, the aliquot is first diluted in PBS1X (as in *i*) and then processed to purify cfDNA, as in *ii*, to check whether the sole dilution of the plasma (performed to reduce plasma viscosity) interferes with the efficiency of cfDNA recovery.

#### EV isolation by ultracentrifugation (UC) and cell-free DNA (cfDNA) extraction

EVs were separated by UC on an Optima MAX-XP ultracentrifuge (Beckman Coulter) equipped with a TLA55 rotor. The plasma samples were filtered (Minisart NML syringe filters, pore size: 0.8 µm; Sartorius) and cleaned from cell debris by two serial centrifugation steps: 2000g for 10min and 10.000g for 20min. EVs were then pelleted at 100.000g for 70min, washed with 1mL of 1xPBS (Gibco), and re-pelleted at 100.000g for 70min as previously reported [5]. The liquid fractions (plasma and washing PBS) recovered from the UC steps were pooled in a separate clean tube and processed for cfDNA isolation as before [3].

*EV isolation by Size Exclusion Chromatography (SEC) (Izon qEV2 columns) and cfDNA extraction:* Plasma samples were filtered (Minisart NML syringe filters, pore size: 0.8 µm; Sartorius) and cleaned from debris by two serial centrifugation steps: 1.500g for 10 min followed by 10.000g for 10 min before loading on a pre-rinsed qEV2/70 nm column (Izon Science LTD, Cat. No. SP4). EVs were collected from 1^st^ to 5^th^ fractions by using an Izon Automatic Fraction Collector (AFC) and concentrated in Amicon Ultra 15 filtering units (Merck Millipore, Cat. No. UFC910024) by centrifugation at 3.000g for 20 min on a 5810R benchtop centrifuge (Eppendorf). The volumes eluted between the 6^th^ and the 20^th^ fractions were collected and processed for cfDNA isolation as before [3].

### EV isolation by Charge-Based (CB) method from cell culture media (CCM)

The SKBR3 and MDA-MB-231 cell lines were cultured in McCoy’s 5A and DMEM medium (26600-023, 11960-044, Gibco), respectively. The media were supplemented with 10% FBS (10270-106, Gibco), 1% Glutamine (25030-024, Gibco), and 1% Pen/Strep (30-002-CI, Corning). Before collecting CCM, the cells were washed two times with 10 mL of PBS 1X (Gibco) and maintained for 24h in regular medium without FBS. The CCM was processed to remove cell debris with two sequential centrifugations (300g x 10 minutes, 2800 rpm x 10 minutes) and filtered (Minisart NML syringe filters, pore size: 0.2 µm; Sartorius). Afterward, the CCM was concentrated with Amicon Ultra centrifugal devices MWCO 100 kDa (Merck) for 10 minutes at 3000g. EVs were isolated from concentrated CCM by the CB method as previously described [1].

### Quantitation of EVs

The size and concentration of the EVs isolated from BrCa patients were quantitated using Tunable Resistive Pulse Sensing (TRPS, qNANO instrument, Izon Science). An average of 2 min recording time was used with nanopores NP250 (A58579, A68549; Izon Science). Voltage was set between 0.30 and 0.40 to achieve and maintain a stable current in the 95-130 nA range, noise between 7-12pA and a linear particle count rate. Calibration was performed using a known concentration of beads CPC200B (mode diameter: 210nm) or CPC400E (mode diameter: 340nm; Izon Science). All the acquisition data were recorded and analyzed by Izon Control Suite v.3 software. Based on instrumentation availability, size distribution profiles and concentration of the EVs isolated from healthy donors’ plasma were obtained using Nanoparticle Tracking Analysis (NTA, NanoSight NS300, Malvern Panalytical). All EV samples were diluted 1:40-1:2000 in 1x Phosphate-Buffered Saline without calcium and magnesium buffer (Gibco) to obtain between 20 and 120 particles /frame. For each sample, 5x 60 s videos were recorded in standard mode with the equipped sCMOS camera (camera level 15-16) and analyzed by NTA 3.4 software (Malvern Panalytical) with a detection threshold ranging from 3 to 5.

### DNA Extraction from EVs

DNA was extracted from EVs by using DNeasy Blood and Tissue kit (Qiagen; cat.69504) by following the manufacturer’s instructions with a few modifications: EVs were resuspended in 200uL of PBS1x before adding 20uL of provided Proteinase K, 4uL of RNAseA (Qiagen; cat. 19101, 100ug/mL) and 200uL of AL buffer (without ethanol). Samples were incubated at 56°C for 10min, mixed with 200uL of 100% ethanol (Sigma-Aldrich) to obtain a homogeneous solution, and pipetted into a DNeasy mini spin column. Afterward, columns were washed with 500uL of AW1 and AW2 buffers and dried at room temperature for 2 min. EV-DNA was eluted by adding 40uL of 10mM Tris-HCl ph8.0 and centrifuging columns at 6.000g for 2min. Measurements of DNA quantity were performed on 10uL of eluted EV-DNA by High Sensitivity dsDNA Qubit Assay (Thermo Fisher Scientific).

### DNA-encapsulating liposomes

DNA-encapsulating liposomes utilized as internal positive control were prepared by the hydration of a thin lipid film. Briefly, 220 μL of 40 mg/mL POPC (1-palmitoyl-2-oleoyl-glycerol-3-phosphocholine; Avanti Polar Lipids) and ten mol% cholesterol (0433-250G, VWR Chemicals) in chloroform (288306, Sigma-Aldrich) were deposited in a 3 mL glass round bottom flask, and the solvent evaporated with a Buchi rotary evaporator. The thin lipid film was kept under vacuum for two hours before DNA encapsulation. Plasmid DNA (1.3 μg, 4Kb) was dissolved in sterile PBS1x (Gibco) and then added to the lipid film and thoroughly vortexed for four min. 5 freeze-thaw cycles were performed to increase encapsulation efficiency. The liposomes were then transferred to 1 mL glass vials and tumbled overnight at RT on a tube rotator unit (10136-084, VWR). After tumbling, the liposomes were extruded through 400 nm Whatman Nuclepore track-etched polycarbonate membranes with an Avanti Mini-Extruder (11 passes) and purified by size exclusion chromatography with Sepharose 4B resin (GE17-0120-01, Cytiva) equilibrated with PBS1x (Gibco). Fractions were collected with a Gilson FC204B multichannel fraction collector. The purified liposomes were utilized within a few hours.

### RNA Extraction from EVs

EV-RNA was isolated by adding TRIzol Reagent (Invitrogen) and chloroform to samples. The RNA-containing phase was separated by centrifugation (10.000 rpm x 15min), mixed with isopropanol (Sigma-Aldrich), and incubated with glycogen (Thermo Fisher Scientific, RNA grade) at −20°C overnight. RNA was precipitated by centrifugation (12.000 rpm x 40min at 4°C) and washed with 70% ethanol by two sequential centrifugation steps (12.000 rpm x 20min; 12.000 rpm x 20min). Dried RNA pellets were further purified with a Single Cell RNA Purification Kit (Norgen; cat.51800). EV-RNA were analyzed on Agilent 2100 Bioanalyzer G2939A coupled with 2100 Expert version 2.6 software.

### Quantitation of cfDNA and bioanalyzer analysis

The size distribution of cfDNA fragment length was analyzed by Agilent High Sensitivity DNA Kit on Agilent 2100 Bioanalyzer G2939A and quantitated by Qubit® dsDNA HS (High Sensitivity) Assay Kit (Thermo Fisher Scientific, Inc.).

### Quantitation of EV-RNA and bioanalyzer analysis

RNA extracted from EVs was analyzed for size distribution and quantitation of fragments length by Agilent RNA 6000 Pico Assay or Agilent Small RNA Assay on Agilent 2100 Bioanalyzer G2939A.

### Digital PCR

Copy Number Determination Assay was performed on cfDNA according to ddPCR Supermix for probes (No dUTP) (Bio-Rad Laboratories Inc.; #1863024). Specifically, 2x ddPCR ddPCR Supermix for probes (No dUTP), 20x target probe (FAM) and 20x reference probe were loaded to 1x as final concentration in a final volume of 22uL. Template (cfDNA) was loaded in a final volume of 6uL without prior digestion with restriction enzymes. Cycling conditions were: 95°C for 10 min, 39 cycles of 94°C for 30 sec and 60°C for 1min followed by 98°C for 10 min and holding at 4°C until droplets reading.

Gene expression Assay was performed on EVs-RNA according to the One-Step RT-ddPCR Advanced Kit for Probes (Bio-Rad Laboratories, Inc.; #1864021). Reaction mix was prepared according with the manufactures’ protocol in a final volume of 22uL. Supermix, 20x target probe (FAM) and 20x reference probe were loaded to 1x as final concentration. Reverse transcriptase was 20U/uL, except in negative control wells. Reverse transcription was performed at 42°C for 60 min and immediately followed by 95°C for 10 min to activate enzyme. Denaturation and Annealing/Extension were 40 cycles of 95°C for 30 sec and 59-60°C for 1 min. enzyme was deactivated at 98°C for 10 min and PCR reactions hold at 4°C until reading. Probes for *ERBB2* (dHsaCP1000116; dHsaCPE5037554), EIF2C1 (dHsaCP2500349), EEF2 (dHsaCPE5050049) were from Bio-Rad Laboratories Inc. Amplification on cfDNA was determined as the ratio between circulating ERBB2 and reference gene EIF2C1 by using a previously reported ddPCR assay for ERBB2 copy number alteration on BrCa tissues [6]. The expression on EV-RNA was calculated as a ratio between the detection of fragments corresponding to ERBB2 and the commonly utilized tissue reference gene EEF2 [7]. PCR reactions were run on T100 thermal cycler (Bio-Rad, Hercules, CA, USA). ddPCR reaction conditions were verified using varied amounts of DNA, RNA and EV-RNA extracted from HER2+ and HER2- cell lines. Reactions without template and /or enzymes were run as internal negative controls. Droplets were generated by QX200 AutoDG Droplet Digital PCR System, and PCR plates were analyzed on a Bio-Rad QX200 droplet reader (Bio-Rad Laboratories, Inc.). Analysis of ddPCR data was analyzed using QuantaSoft Software, version 1.7 (Bio-Rad Laboratories, Inc.).

### EV protein isolation and Western blot analysis

EV proteins were extracted and analyzed by western blotting as previously reported [8] with few modifications: total EV proteins were loaded on 4-15% mini – PROTEAN TGX stain-free gels (Bio-Rad), transferred on a trans-blot turbo system (Bio-Rad) and visualized on a UVITEC imaging system (UVItec Ltd, Cambridge). Primary antibodies used were rabbit anti -CD9 (#D801A; Cell signaling Technology; 1/500); rabbit anti – CD 9 (# MA5-33125; Invitrogen; 1/500); mouse anti – Apolipoprotein A1 (#5F4; Cell Signaling Technology; 1/250); rabbit anti– Albumin (#4929; Cell Signaling Technology; 1/1000), mouse anti-Tsg-101 (ab83, Abcam, 1/1000), rabbit anti-Flotillin 1 (#18634, Cell Signaling Technology; 1/1000), rabbit anti-Flotillin 2 (#3436, Cell Signaling Technology; 1/1000), rabbit anti-CD44 (ab157107, Abcam, 1/500), rabbit anti-EpCam (#D1B3, Cell Signaling Technology; 1/500), rabbit anti-HER2/ErbB2 (#29D8, Cell Signaling Technology; 1/1000).

### Imaging Flow Cytometry

Isolated EV samples (10^10^-10^11^ particles/mL) were resuspended in 1x Phosphate-Buffered Saline (1xPBS) (Gibco) and incubated overnight at 4C with HER2/ErbB2 (29D8) – PE-conjugated antibody (1:50) or concentration-matched Rabbit (DA1E) mAb IgG XP® Isotype Control (PE Conjugate) (Cell Signaling Technology). EVs were then stained with one volume of CellMask plasma membrane stains (C10046, Life Technologies) diluted in 1xPBS (1:5000) for 15 minutes at room temperature, in the dark. Before acquisition on Amnis ImageStream X MkII, the stained samples were diluted with 1xPBS to 10^8^ EVs / mL to avoid coincidence.

All samples were analyzed on an Amnis ImageStream X MkII (Luminex). All data sets were acquired with a 60x objective on High Gain mode using INSPIRE® instrument acquisition software (Luminex). Fluidics was set to low speed; sensitivity was set to high resolution.

Samples were acquired with all lasers run at maximal power (488nm:200mW, 642nm:150mW, 758nm:70mW). Sub-micron beads (F13839, Life Technologies) were run after instrument initialization as an internal standard. Double-stained EV samples from BrCa patients were loaded on tubes and acquired for the same time (3 minutes). To avoid the carryover of fluorophores and dyes during the acquisition, washes with deionized water (0.1 μm filtered) were performed between samples. The following control samples were run before double stained samples as internal controls: i) 1x Phosphate-Buffered Saline (PBS) without calcium and magnesium buffer (Gibco) only; ii) unstained EVs; iii) staining solutions (1x PBS plus CellMask or PE-conjugated antibodies).

### Multiplex EV surface marker analysis

Isolated EVs (5*10^8^ - 10^9^ particles counted by NTA) were analyzed for surface protein expression using the MACSPlex Exosome kit (Miltenyi Biotech; no. 130-108-813) by following the manufacturers’ protocol tube for overnight capture. Detection was performed by incubating samples with 15μL of the provided MACSPlex Exosome Detection Reagent cocktail for 2 hours at room temperature. All samples were run as triplicates, and PBS (without EVs) was included as blank control. Data were acquired on a BD FACSCanto flow cytometer (BD Biosciences) and analyzed with FlowJo v10 Software (BD Life Sciences). Normalized and background-corrected MFI values were used for sample comparison.

### *In situ* analyses of breast cancer patient clinical cohort

Fluorescence in situ hybridization (FISH) assay for ERBB2 was performed using the ZytoLight CEN17/SPEC ERBB2 Dual Color Probe (ZytoVision) by following the ZytoLIGHT Implementation KIT for in situ hybridization. Slides were analyzed on a Zeiss Axio-Imager 2 according to ASCO-CAP guidelines [9]. Immunohistochemistry staining was performed as

part of the clinical laboratory workflow per guidelines. All *in situ* analyses were performed at the Unit of Surgical Pathology of the Santa Chiara Hospital in Trento, Italy.

### DNA whole-exome sequencing

For library preparation (SeqCap EZ HyperCap Workflow version 2.3; Roche), we utilized 20-50ng of cfDNA and 100ng of matched germline DNA (gDNA) sonicated to reduce the size to 180-220bp (Covaris M220). Libraries were sequenced on Illumina HiSeq2500 platform by the Next Generation Sequencing Facility at the University of Trento (Italy) with a Paired-End, 100bp protocol with a mean coverage of 537x (min=282x, max=731x) for cfDNA and of 96x for gDNA (min=89x, max=109x) (**Supplementary Table 4**). Fastq files were controlled for quality by fastqc (www.bioinformatics.babraham.ac.uk) and trimmed by Trimmomatic [10] (SLIDINGWINDOW:4:15 MINLEN:36). Alignment was performed using BWA [11] with default parameters. Duplicate removal, realignment around indels, and base quality recalibration were performed with MarkDuplicates, RealignerTargetCreator, IndelRealigner, BaseRecalibrator and ApplyBQSR from gatk4 pipeline [12]. The genetic match of cfDNA and control gDNA was verified by SPIA [13]. CNVkit [14] was used for copy number aberration (CNA) detection via segmentation, and Log2 values of cfDNA over control were corrected for purity and ploidy using ClonetV2 [15]. Results were visualized using IGV [16] and custom R scripts (libraries: circlize [17]). See also **Supplementary Table 2**.

### RNA-Seq

To generate libraries for RNA-Seq, the purified EVs-RNA were processed with SMART-Seq® Stranded Kit (Takara Bio USA, Inc.). Libraries were sequenced on the Illumina HiSeq2500 platform by the Next Generation Sequencing Facility at the University of Trento (Italy) with a Single-End, 100bp Single-End protocol generating an average of 88M reads (min=56, max=97) (**Supplementary Table 5**). Fastq files were controlled for quality and trimmed using fastp [18]. Reads were aligned against the human genome (hg38) using STAR [19] (custom parameters, outFilterMatchNminOverLread 0.3, outFilterScoreMinOverLread 0.3).

Breast cancer tissue data [20] were downloaded from CBioPortal (https://www.cbioportal.org/). Capped relative linear copy-number values were used for DNA. RNA Seq V2 RSEM was used for RNA. Clinical data annotation includes immunohistochemistry evaluation of HER2.

### HER2 positivity classification of plasma and tissue samples

For HER2 positivity classification of BrCa study patients, we used ddPCR data for all liquid biopsy samples (the highest DNA and RNA ddPCR values of healthy donor samples were used as lower thresholds for DNA and RNA, respectively) and sequencing data for the TCGA tissue samples (DNA amplification threshold was set at 2.6 copies; RNA levels for samples without DNA amplification (DNA copies < 2.6) were considered and the 75% percentile of the distribution was set as lower threshold).

For each sample (either plasma or tissue), we then considered the following classes of HER2+ positivity; *Only RNA*: RNA but not DNA signal is higher than the corresponding threshold; *Only DNA*: DNA but not RNA data is higher than the corresponding threshold; *Combo AND*: both DNA and RNA data are above the corresponding thresholds; *Combo OR*: any of DNA or RNA data is higher than the corresponding threshold.

For classificatory accuracy estimation, precision is defined as:

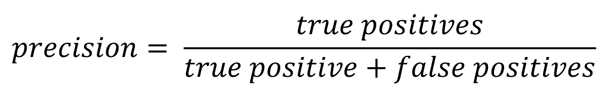

Recall (i.e., sensitivity) is defined as:

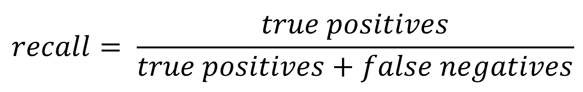

Specificity is defined as:

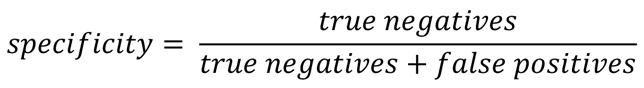

F1 score is defined as:

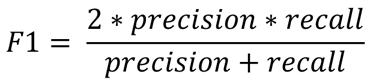

All performance measures of plasma and tissue-based HER2 positivity are shown in **Supplementary Table 5.**

